# Glutathione S-transferase Pi (Gstp) Proteins Regulate Neuritogenesis in the Developing Cerebral Cortex

**DOI:** 10.1101/2020.05.21.103036

**Authors:** Xiaonan Liu, Sara M. Blazejewski, Sarah A. Bennison, Kazuhito Toyo-oka

**Affiliations:** Department of Pharmacology and Physiology, Drexel University College of Medicine Department of Neurobiology and Anatomy, Drexel University College of Medicine Philadelphia, PA 19129 USA

## Abstract

GSTP proteins are metabolic enzymes involved in removal of oxidative stress and intracellular signaling and also have inhibitory effects on JNK activity. However, the functions of Gstp proteins in the developing brain are unknown. In mice, there are three Gstp proteins, Gstp1, 2 and 3, while there is only one GSTP in humans. By RT-PCR analysis, we found that Gstp1 was expressed beginning at E15.5 in the cortex, but Gstp2 and 3 started expressing at E18.5. Gstp 1 and 2 knockdown caused decreased neurite number in cortical neurons, implicating them in neurite initiation. Using *in utero* electroporation to knockdown Gstp1 and 2 in layer 2/3 pyramidal neurons *in vivo*, we found abnormal swelling of the apical dendrite at P3 and reduced neurite number at P15. Using time-lapse live imaging, we found that the apical dendrite orientation was skewed compared to the control, but these defects were ameliorated. Overexpression of Gstp 1 or 2 resulted in changes in neurite length, suggesting a role in neurite elongation. We explored the molecular mechanism and found that JNK inhibition rescued reduced neurite number caused by Gstp knockdown, indicating that Gstp regulates neurite formation through JNK signaling. Thus, we found novel functions of Gstp proteins in neurite initiation during cortical development. Furthermore, the overexpression experiments suggest different functions of Gstp1 and 2 in neurite elongation. Since previous studies have shown the potential implication of Gstp in Autism Spectrum Disorder, our findings will attract more clinical interests in Gstp proteins in neurodevelopmental disorders.

**Significance:** Neurite formation, including neurite initiation and elongation, is the first step of generating polarized neuronal morphology in developing neurons, and thus is essential for establishing a neuronal network. Therefore, it is crucial to understand the mechanisms of neurite formation. Limited studies have been performed to clarify the mechanisms of neurite formation, especially neurite initiation. In this present study, we report a novel, essential role of Gstp in neurite initiation in mouse cortical neurons *in vitro* and *in vivo*. We also found that Gstp regulates neurite formation via JNK signaling pathways. These findings not only provide novel functions of Gstp proteins in neuritogenesis during cortical development but also help us to understand the complexity of neurite formation.

## Introduction

Neurite formation includes neurite initiation and neurite elongation, and is the process where a neuron starts to gain morphological polarity. It is an extremely complicated process where a lot of internal and external factors are involved in both steps (Drubin et al., 1985; Perron and Bixby, 1999). Neurite formation is fundamental for the development of the central nervous system, as it creates structural basis for neuronal connection, communication, and plasticity (Reese and Drapeau, 1998). In particular, neurite initiation is the cornerstone of neurite formation which neurite number primarily relies on (Schaefer et al., 2008; Harrill et al., 2010). Studying this process can help us understand the etiology of neurodevelopmental disorders, for example, autism spectrum disorder (ASD) and attention-deficit/hyperactivity disorder (ADHD), because recent studies have implicated defects of neuritogenesis in these disorders (Won et al., 2011; Bakos et al., 2015). Moreover, the model system used for the mechanistic analysis of neurite formation is shared with the study of neurite and axonal degeneration.

GSTP belongs to the glutathione S-transferase (GST) family (Mannervik et al., 1985). GSTs are enzymes that catalyze the conjugation of glutathione to molecules, and therefore help further metabolize and excrete molecules from the cell. This reaction is critical because it is involved in detoxification and removal of oxidative stress (Goto et al., 2009; Aaker et al., 2016). There is one GSTP protein in humans, GSTP1. The human *GSTP1* gene is encoded in chromosome 11q13 in the genome. It has been reported that a GSTP1 single nucleotide polymorphism (SNP) is associated with the neurological disorder, Tourette syndrome, which share some similar symptoms with ASD (Darrow et al., 2017). A SNP on the promoter region of the *GSTP1* gene has a significant association with this disorder (Shen et al., 2014). In mice, there are three Gstp genes, *Gstp1, Gstp2* and *Gstp3*, which are encoded by three different, but adjacent regions on chromosome 19 (Xiang et al., 2014). Previous research showed Gstp1 and 2 are ubiquitously expressed during embryonic stages in the central nervous system and throughout the mouse body except for the uterus (Knight et al., 2007). Gstp3 was discovered more recently, and a limited number of studies have been done in terms of its expression and functions. Besides the enzymatic activity important for the detoxification of oxidative stress, GSTP1 is also involved in cellular signaling and cell proliferation (Zhang et al., 2014). Studies have shown that GSTP1 can inhibit the activation of several kinases, including JNK1(MAPK8) and Cdk5 (Adler et al., 1999; Wang et al., 2001; Sun et al., 2011). Thus, GSTP proteins have multiple functions and are essential for cellular signaling important for various types of cellular events.

JNKs are kinases essential for cell proliferation and apoptosis. There are three JNK-encoding genes, and each of them can be alternatively spliced to form several variants (Gupta et al., 1996). It has been implicated that the C-terminus of GSTP1 directly interacts with the C-terminus of JNK1 (Monaco et al., 1999; Wang et al., 2001). This interaction leads to an inhibitory effect on JNK1 activation, which affects the cellular signaling cascade mediated by JNK1 activation. Also, GSTP1 binds to JNK2 and inhibits JNK2 activity (Thévenin et al., 2011). While there is no direct evidence showing the interaction between GSTP1 and JNK3, JNK3 has high homology to JNK1 suggesting that GSTP1 may also interact with JNK3 in a similar fashion as JNK1 (Sun et al., 2011). JNK signaling pathways are most notable for reacting to oxidative stress and inducing apoptosis (Shen and Liu, 2006). However, there is increasing evidence showing that JNK has non-apoptotic functions in neurons and is necessary for neuronal development (Eom et al., 2005; Seow et al., 2013). It has been shown that JNK proteins play multiple roles in neurite outgrowth (Bennison et al., 2020). JNK1 is important for neurite elongation, and JNK2 and 3 are involved in both neurite initiation and elongation (Barnat et al., 2010). Notably, JNK proteins are involved in cytoskeletal organization via a variety of factors, including microtubule-associated protein 2 (MAP2) and paxillin (Yamauchi et al., 2006; Komulainen et al., 2014). This is critical for neuritogenesis because the organization of cytoskeletal components, actin and microtubules, is key for neurite initiation and elongation, since they build tension and push the membrane outward to break the spherical shape, as well as push the neurite to grow in the later stages of neurite formation (Flynn, 2013).

Thus, the previous studies about the link between JNKs and Gstp proteins implicates Gstp as an upstream regulator of JNKs in neurite formation. However, little has been studied about Gstp proteins in the developing cerebral cortex. In this study, by knocking down Gstp1 and 2 together in mouse primary neurons, we found that Gstp1 and 2 are involved in the formation of the correct number of neurites, suggesting their importance in neurite initiation. *In vivo* knockdown by *in utero* electroporation in the developing cerebral cortex showed defects in orientation of the apical dendrite at P3 and in neurite initiation of basal dendrites at P15. *Ex vivo* time-lapse live imaging of the P0 brain showed that the morphology of Gstp1/2-knockdown neurons dramatically changed with a disrupted angle of the apical dendrite as it emerged from the soma, suggesting that Gstp1 and 2 are important for correct apical dendrite orientation. Overexpression of each Gstp protein in primary cortical neurons revealed that Gstp1 overexpression caused decreased length of the shorter neurites, which will likely become dendrites, while Gstp2 overexpression caused a decrease in length of the longest neurite, which will likely become the axon. By applying a global JNK inhibitor, which inhibits JNK1, 2 and 3, to Gstp-deficient neurons, we found that the inhibition of JNKs’ activity rescued the defects in neurite initiation caused by Gstp knockdown, indicating the importance of the Gstp/JNK signaling pathway in neurite initiation. Thus, our results provide the first evidence that Gstp1 and 2 are essential regulators of neuritogenesis, especially during neurite initiation via the JNK signaling pathway in the developing cortex.

## Materials and methods

### Mice

CD-1 mice used for IUE and primary cortical neuron culture were purchased from the Charles River (Wilmington, MA). All animals were maintained in accordance with the guidelines of the Drexel University Institutional Animal Care and Use Committee. Females and males were used for *in utero* electroporation and primary cortical neuron culture.

### Plasmids and chemicals

Myc and DDK (FLAG)-tagged mouse Gstp1, Gstp2 and Gstp3 cDNA clones in the pCMV6-Entry vector were purchased from Origene (MR202273, MR202254, MR202253, Rockville, MD). shRNA for the knockdown of both Gstp1and Gstp2, but not Gstp3, was designed by utilizing the web tools. They are siRNA Wizard Software (InvivoGen), BLOCK-iT RNAi Designer (ThermoFisher Scientific), and GPP Web Portal (Broad Institute). Target sequence, GGAGGTGGTTACCATAGAT, was cloned into pSCV2-Venus plasmid (Hand and Polleux, 2011; Toyo-oka et al., 2014). ACTACCGTTGTTATAGGTG was used as a scramble shRNA. SP600125 (JNK inhibitor), and Roscovitine (Cdk5 inhibitor) were obtained from APExBIO (Houston, Texas).

### Antibodies

The following primary antibodies were used: anti-GSTP1 (Rabbit, Proteintech, 15902-1-AP), anti-GAPDH (Mouse, Proteintech, 60004-1-Ig), anti-DYKDDDDK (FLAG) Tag (Rat, Biolegend, clone L5, 637301), anti-Brn2 (Rabbit, Proteintech, 14596-1-AP). Anti-βIII-tubulin (mouse, Thermo Scientific, clone 2G10, MA1-118). The following secondary antibodies were used: HRP-conjugated Goat anti-Rabbit IgG (Jackson ImmunoResearch Laboratories, 111-035-003), HRP-conjugated Goat anti-mouse IgG (Proteintech, SA00001-1), HRP-conjugated Donkey anti-Rat IgG (Jackson ImmunoResearch Laboratories, 712-035-153), FITC-conjugated Donkey-anti-Rabbit IgG (Jackson ImmunoResearch Laboratories, 711-096-152), TRITC-conjugated Donkey-anti-Rabbit IgG (Jackson ImmunoResearch Laboratories, 711-025-152), FITC-conjugated Donkey-anti-mouse IgG (Jackson ImmunoResearch Laboratories, 715-096-151).

### Validation of shRNA knockdown efficiency

HEK-293 cells were co-transfected with the plasmids coding the scramble shRNA or the Gstp1/2 shRNA and Myc and DDK (FLAG)-tagged mouse Gstp1, Gstp2, and Gstp3 plasmids. The cells were cultured for 48 hours at 37°C with 95% air/5% CO_2,_ and the protein lysates were prepared. The protein samples were separated on 12% SDS-PAGE gel. Knockdown efficiency was analyzed by quantifying the expression level of FLAG-tagged Gstp proteins after blotting with anti-FLAG antibody. GAPDH was used as a loading control. To analyze the KD efficiency on endogenous Gstp proteins, the plasmids coding scramble or Gstp shRNA were transfected into N2a cells and analyzed the efficiency with anti-GSTP1 antibody after 48 hours as described above.

### In Utero Electroporation (IUE)

Timed pregnant mice were obtained by the set-up of the mating in the animal facility. *In utero* electroporation (IUE) was performed as previously described (Cornell et al., 2016). Briefly, after pregnant dams were anesthetized, the uterine horns were exposed, and one to two microliters of plasmid mixed with 0.1% Fast Green were injected into the lateral ventricle of E15.5 embryo brains by pulled-glass micropipette. The concentration of the plasmid DNA was 1-2μg/μl. Three 32V electric pulses were applied into the embryonic brain by tweezers electrode using the electroporator (CUY21, Nepa GENE). The uterine horns were returned into the abdomen, and pups were allowed to recover and mature. The brains were dissected out at P0 for time-lapse live imaging and P3 and P15 for morphological analysis.

### Primary Cortical Neuron Culture

Cortical neurons were prepared from mouse embryos at E15 as previously described (Pischedda et al., 2018). Briefly, the dam was euthanized, and embryos were quickly removed from the pregnant mouse. Then, embryos were placed in 1X Ca^2+^/Mg^2+^-free Dulbecco’s PBS (D-PBS, Genesee Sci). After the cerebral cortices were dissected out, they were treated with 0.01% Trypsin in D-PBS for 5 minutes at room temperature, and then the trypsin is neutralized by adding 100μl of 50mg/ml bovine serum albumin. The dissociated neurons were seeded onto the dish coated with 100 ng/ml poly-D-lysine and 100ng/ml laminin with Neurobasal medium supplemented with 1% penicillin/streptomycin (Corning), 1% GlutaMAX (Gibco), and 1X B-27 (Thermo Fischer Scientific). After 48 hours, cells were re-plated onto glass coverslips coated with poly-D-lysine and laminin and cultured for an additional 48 hours, and then fixed with 4% paraformaldehyde/Phosphate-buffered saline.

### Transfection

Transfection in primary cortical neurons was performed using Amaxa Nucleofector II (Lonza) with the Ingenio electroporation kit (Mirus Bio). The concentration of plasmids we used was 10μg with 100 μl of Ingenio electroporation solution. Five million neurons were used for transfection per experimental group.

### RNA isolation and RT-PCR

The cerebral cortices were dissected from the cerebral cortex at E15.5, E18.5, P0, P5, and P15, and the total RNA was prepared using Trizol reagent (Thermo Fisher Scientific). The quality of RNA was confirmed by the value of 260nm/280nm and the clear appearance of 18S and 28S rRNAs on the agarose gel. To create cDNA for PCR, RNA was reversely transcribed using the MMLV reverse transcriptase (Thermo Fisher Scientific) and Oligo (dT) primer (Promega), and the following heat cycle was used for the reverse transcription: 25°C 10 minutes, 37 °C 60 minutes, 70 °C 10 minutes. Specific Gstp1, 2, and 3 primers were designed using the information about their DNA sequence. Gstp1 primers; forward primer: GGCAAATATGTCACCCTCATCTACACC, reverse primer: CCTTGATCTTGGGCCGGGCAC, Gstp2 primers; forward primer: CGGCAAATATGGCACCATGATCTACAGA, reverse primer: CCTTGATCTTGGGCCGGGCAC, Gstp3 primers; forward primer: CCTTACACCATCGTCGTCTATTTCCCTTCC, reverse primer: GATACTGCCGGGCAATGCGTCTG. PCR was performed using these specific primers with the following PCR heat cycle condition: 94 °C 5 min and 73 °C s 30 sec. This cycle was repeated 42 cycles. The product sizes are 271, 272 and 319 bp for Gstp1, 2, and 3, respectively.

### Histology

Brains were dissected out at P3 and P15 and fixed with 4% paraformaldehyde/Phosphate-buffered saline at 4°C overnight. Fixed samples were cryoprotected by 25% sucrose/Phosphate buffered saline for 48 hours at 4°C, and then embedded with the O.C.T. compound (Sakura). Cryosections (60 μM thickness) were cut by cryostat (Microm HM 505 N) and air-dried. Sections were washed three times with Tris-buffered saline (TBS) before use. All brain sections were stained with 4’, 6-Diamidino-2-phenylindole, Dihydrochloride (DAPI, 600nM) and embedded with 90% glycerol/Phosphate buffered saline.

### Neuromorphological Analysis

To analyze the neuronal morphology at P3 and P15 and primary neurons, Fiji software was used. Z-projection images were created from z-stack data collected by the confocal microscope (TCS SP2, Leica). The Fiji plug-in Simple Neurite Tracer was used to measure the length of the neurites extended from the surface of soma.

To analyze the apical dendrite orientation at P3, the angle was measured using the Fiji software angle tool. A straight line perpendicular to the pial surface of the brain slice was used as a reference for the angle. Absolute values of the angles measured were used for statistics.

### Sholl analysis

Neurite branching pattern was quantified using Sholl analysis in Fiji software as previously described (Cornell et al., 2016). The center of soma was used as a reference, and the radius was set to 200 or 250 μm with 5 μm interval. From these parameters, the number of intersections at each radius was quantified and plotted using Prism7 (GraphPad).

### Ex Vivo Live Imaging on Brain Slices

Brain slices were collected as previously described with a slight modification (Cornell et al., 2016). Briefly, Brains were dissected out from P0 embryos and placed in an ice-cold high sucrose artificial cerebral spinal fluid (ACSF) solution. The brains were embedded in 4% low-melting agarose in the ACSF solution. Coronal cortical slices were sectioned (300 µm) with a vibrating microtome (VTS1000 Leica Microsystems) in ACSF. Slices were incubated for 60 minutes at 37°C in DMEM/F-12 imaging media supplemented with 10% fetal bovine serum (FBS) for recovery. Slices were gently transferred into a 35 mm Petri dish (Nunc) for imaging. The brain slices were covered by 80% collagen I (Gibco, Fisher Scientific) neutralized with 0.25 N sodium hydroxides in PBS. Imaging was performed on an upright confocal laser-scanning microscope (TCS SP2 VIS/405, Leica) with a 20X HCX APO L waster-dipping objective (NA 0.5). During imaging, slices were incubated in DMEM/F-12 imaging media supplied with 10% FBS without phenol red and incubated at 37°C overnight in stage top chamber incubator (DH-40iL, Warner Instruments). Confocal stack images were taken at 10-minute intervals for up to 10 hours. The Z-projections were created using Fuji for each time point, and the z-projections of all time points were used to make a video. The neurite length, velocity, and angle were measured and analyzed using Fuji software and plotted using Prism7 (GraphPad).

### Statistical Analysis

Quantitative data were subjected to statistical analysis using Prism 7 (GraphPad). The data were analyzed by two-tailed unpaired t-tests, one-way ANOVA with Dunnett’s multiple comparisons, one-way ANOVA, or two-way ANOVA with Tukey’s multiple comparisons if appropriate. Values represented as mean ± SEM. Results were deemed statistically significant if the p value was <0.05. *, **, *** and **** indicate p <0.05, p <0.01, p<0.001 and p <0.0001, respectively.

## Results

### Gstp proteins are expressed during cortical development, and their polarized distribution was observed during neurite formation

A previous report showed that Gstp1, 2, and 3 had different expression patterns in the mouse brain (Visel et al., 2004; Diez-Roux et al., 2011). However, as far as we know, there are no specific antibodies for Gstp1, 2, and 3 available commercially. Therefore, we used the anti-GSTP1 antibody to detect the expression level of Gstp proteins. First, we tested the specificity of the antibody against each Gstp (Figure 1A). We overexpressed FLAG-tagged Gstp1, Gstp2, or Gstp3 in HEK-293T cells respectively, and the protein lysates from each group were tested by Western blot. Anti-GSTP1 antibody detected all three Gstp proteins. Using protein lysates from the cerebral cortex at E13.5, E15.5, E17.5, and P0, we tested the expression levels of Gstp proteins during the development of the cerebral cortex and found that Gstp proteins were expressed throughout all tested stages of cortical development (Figures 1B and C).

**Figure 1.**
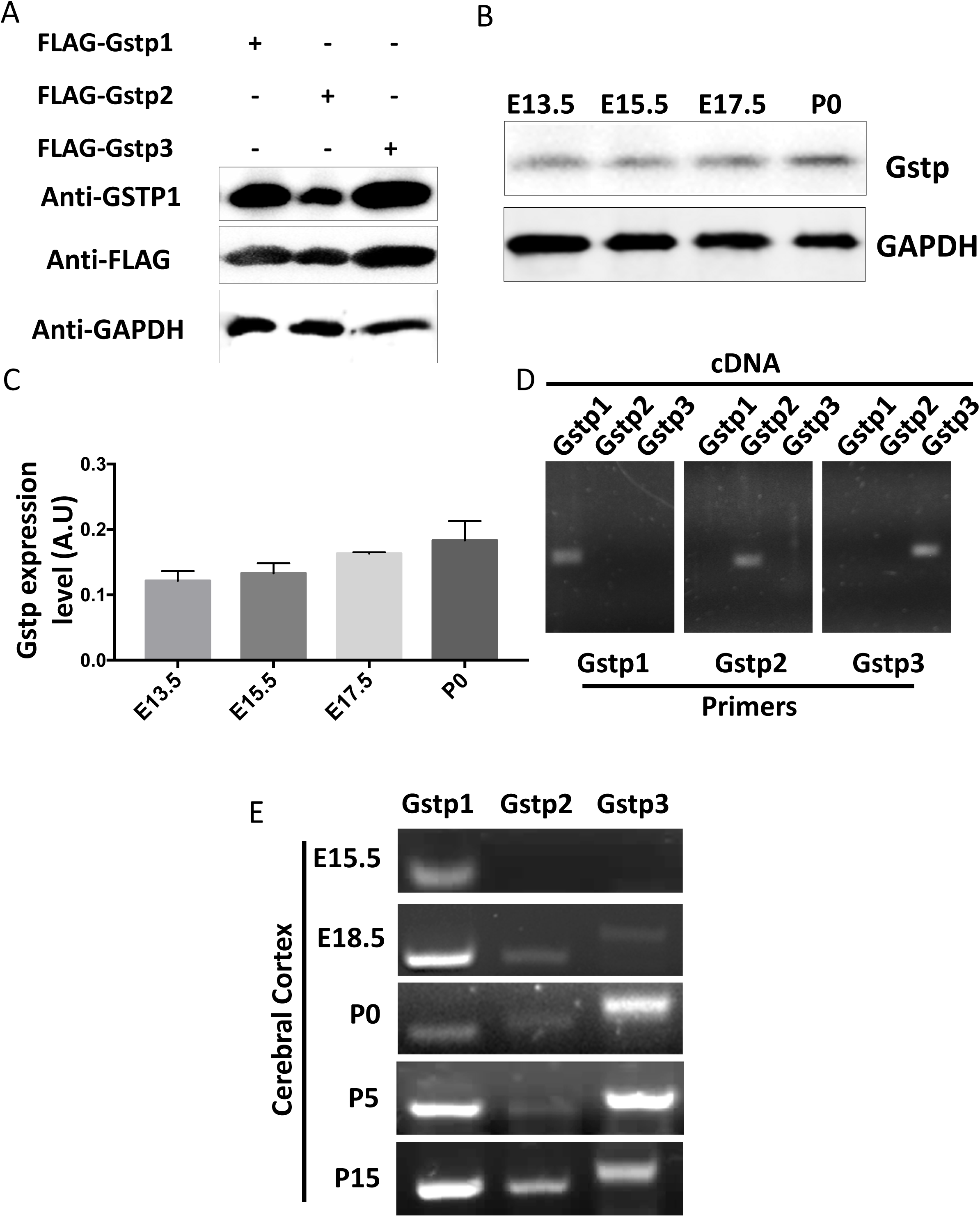
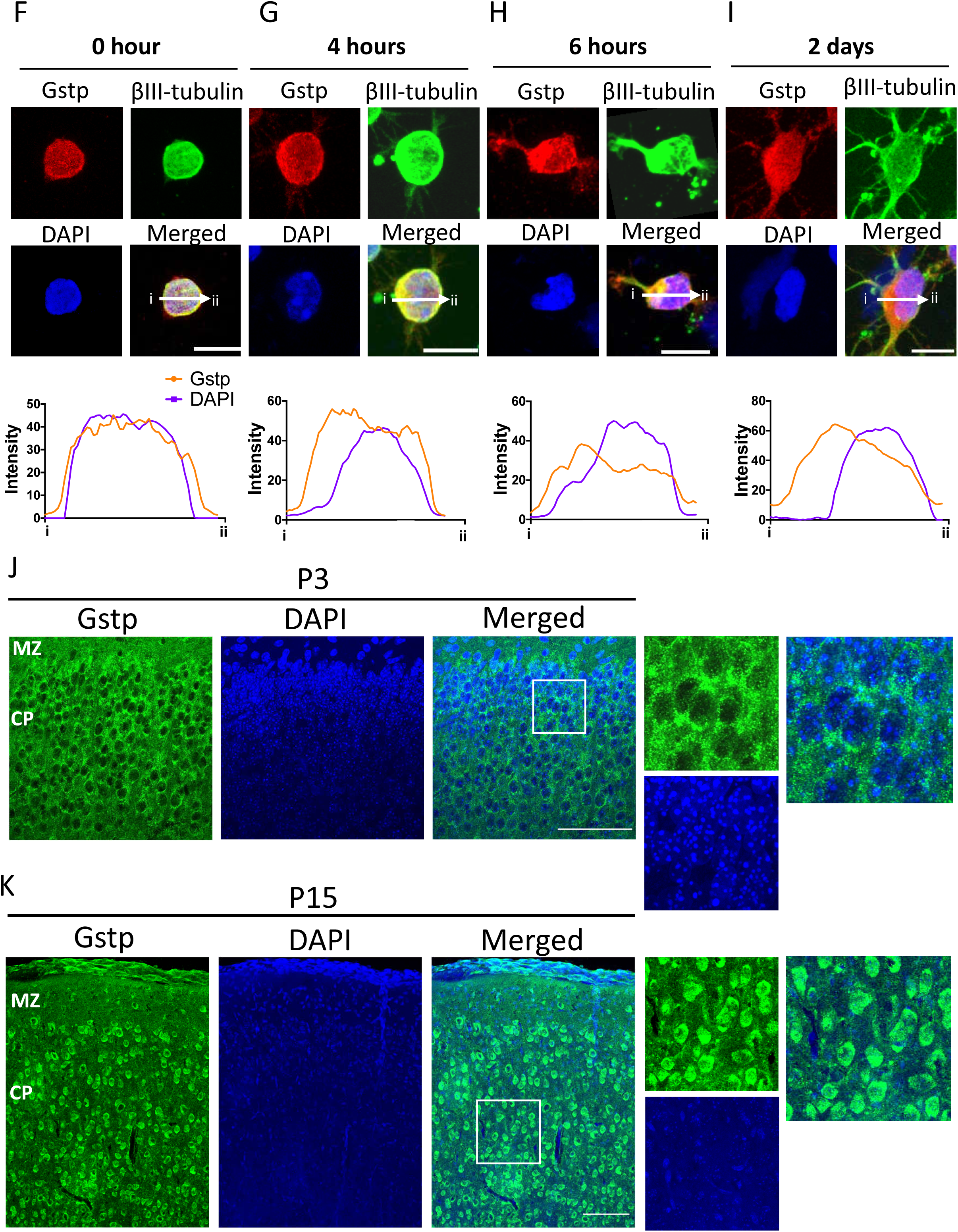
Gstp proteins strongly express inside the soma of mouse cortical neurons during cortical development. **A.** GSTP1 antibody recognizes all three mouse Gstp proteins, Gstp1, 2, and 3. **B.** Gstp proteins express in the developing cortex at E13.5, E15.5, E17.5, and P0. **C.** Quantification of western blot data of Gstp expression in the developing cortex normalized to GAPDH. **D**. Design of the specific primer sets for each *Gstp* gene. RT-PCR revealed that the specific primer set for *Gstp1, Gstp2 and Gstp3* specifically amplified the *Gstp1, Gstp2 and Gstp3*. **E.** mRNA expression of each Gstp mRNA in the developing cortex at E15, E18, P0, P5, and P15. **F-I.** Top photos: Immunostaining of Gstp proteins in primary differentiating cortical neurons. Non-polarized primary cortical neuron (0 hr after plating on a dish, F), Early neurite initiation stage (4 hrs, G), Late neurite initiation stage (6 hrs, H) and Neurite extension stage (2 days, I). Arrows from label “i” to “ii” indicate the regions where the signal intensity was measured. Bottom graphs: Quantification of the signal intensity of DAPI and Gstp crossing the soma from i to ii. Gstp proteins express both in the cytoplasm and the nucleus. Scale bar, 10 μm. **J and K.** Immunohistochemical analysis about the expression of Gstp protein in the developing cortex at P3 (J) and P15 (K).Left panels: low magnification, Right panels: high magnification. Gstp proteins strongly express in the soma and weakly in the proximal part of dendrites. Scale bar, 100 μm.

Since the antibody recognizes all mouse Gstp isoforms, we created specific primer sets for each Gstp mRNA to further examine the expression of each Gstp mRNA in the developing cortex (Figure 1D). Using the plasmids coding Gstp1, 2, and 3 and the specific primers, we performed PCR and confirmed that each primer set is specific for each Gstp gene. Next, we tested the expression pattern of each Gstp mRNA in the developing cortex by RT-PCR and found that Gstp1 started expressing at E15.5 and remained expressed throughout all the time points from E15.5 to P15 (Figure 1E). Gstp2 and 3 started expressing at E18.5, and their expression continued until at least P15. Thus, these experiments suggest that Gstp1 is the main Gstp involved in early cortical development in the embryonic brain.

To determine the cellular localization of Gstp proteins, we conducted immunostaining using the anti-GSTP1 antibody in primary cortical neurons at four different stages of neurite formation, 0 hours after plating (non-polarized stage, Figure 1F), 4 hours (early neurite initiation, Figure 1G), 6 hours (late neurite initiation, Figure 1H), and 2 days (neurite extension, Figure 1I). In non-polarized stage, Gstp was ubiquitously expressed in the cytoplasm and the nucleus (Figure 1F). In the early and late neurite initiation, Gstp expression was observed in the cytoplasm and the nucleus, but concentrated accumulations were observed in the cytoplasm (Figure 1G and H). During neurite extension, Gstp protein was ubiquitously expressed in the cytoplasm, neurites, and the nucleus with less degree (Figure 1H).

Immunohistochemical analysis revealed that Gstp proteins were expressed at the cortical plate (CP) at P3 (Figure 1J) and P15 (Figure 1K). Gstp proteins were strongly expressed in the soma, but weakly or at minimum in the axon and dendrites both at P3 and P15 (Figures 1J and K, right panels).

Thus, Gstp proteins, especially Gstp1, are expressed in the developing cortex. Their polarized expression in the cytoplasm suggests a role for Gstp proteins in neurite formation during cortical development.

### Knockdown of Gstp1 and 2 caused decreased neurite number from the soma and defects in the branching of neurites of cortical neurons

We found that Gstp proteins are expressed in the developing cortex (Figure 1), while their roles in cortical neurons have not been clarified. In mouse, there are three *Gstp* genes encoding Gstp1, 2, and 3, while humans have only one, *GSTP1*. We aligned human GSTP1 and mouse Gstp1, 2, and 3 nucleotide and protein sequences and found that mouse Gstp1 has the highest homology to human GSTP1 with 83.25% identity in nucleotide sequence and 85.24% identity in amino acid sequence (Tables 1 and 2). Mouse Gstp1 and 2 show 98.1% and 97.14% identity in nucleotide and an amino acid sequence, respectively, while mouse Gstp3 shares less similarity with Gstp1 and Gstp2 (71.56% and 71.56% in nucleotide and 70% and 70.48% in amino acid). Therefore, we designed the shRNA in order to specifically knock down both Gstp1 and 2 at the same time, but not Gstp3, by using the website-based siRNA sequence prediction tools (See the Materials and Methods for more details). We confirmed the specificity of the shRNA using HEK-293 cells overexpressing the plasmids coding FLAG-tagged Gstp1, 2, and 3 and scramble shRNA or Gstp shRNA (Figure 2A). The knockdown efficiency was approximately 85% and 95% to Gstp1 and Gstp2, respectively. To analyze the knockdown efficiency of endogenous Gstp proteins, we used a mouse neuroblastoma cell line, N-2a cells. The knockdown efficiency of the Gstp1/2 shRNA was 41% in the endogenous Gstp proteins compared to the cells transfected with plasmid coding the scramble shRNA (Figure 2B). This could be caused because N-2a cells express all Gstp proteins. To confirm this, we performed RT-PCR in N-2a cells and found that N-2a cells expressed Gstp1 and 3, but not Gstp2 (Figure 2C). Thus, we determined that the shRNA we designed was specific to Gstp1 and 2.

**Table 1.** Homology of cDNA sequence among mouse Gstp1, 2 and 3 and human GSTP1

|  |  | *Gstp1* | *Gstp2* | *Gstp3* | *GSTP1* |
| --- | --- | --- | --- | --- | --- |
| **Human** | ***GSTP1*(633 nt)**<br>**NM_000852.3** | 83.25 | 82.46 | 75.2 | 100 |
| **Mouse** | ***Gstp1*(633 nt)**<br>**NM_013541.1** | 100 | 98.1 | 71.56 | 83.25 |
|  | ***Gstp2*(633 nt)**<br>**NM_181796.2** | 98.1 | 100 | 71.56 | 82.46 |
|  | ***Gstp3*(633 nt)**<br>**NM_144869.3** | 71.56 | 71.56 | 100 | 75.2 |

**Table 2.** Homology of amino acid sequence among mouse Gstp1, 2 and 3 and human GSTP1.

|  |  | **GSTP1** | **Gstp1** | **Gstp2** | **Gstp3**<br>**isoform 1** | **Gstp3**<br>**isoform 2** |
| --- | --- | --- | --- | --- | --- | --- |
| **Human** | **GSTP1 (210 aa)**<br>**CAG29357.1** | 100 | 85.24 | 83.81 | 70.48 | 69.95 |
| **Mouse** | **Gstp1 (210 aa)**<br>**NP_038569.1** | 85.24 | 100 | 97.14 | 70.00 | 68.39 |
|  | **Gstp2 (210 aa)**<br>**NP_861461.1** | 83.81 | 97.14 | 100 | 70.48 | 68.91 |
|  | **Gstp3 ispform1 (210 aa)**<br>**NP_659118.1** | 70.48 | 70 | 70.48 | 100 | 100 |

**Figure 2.**
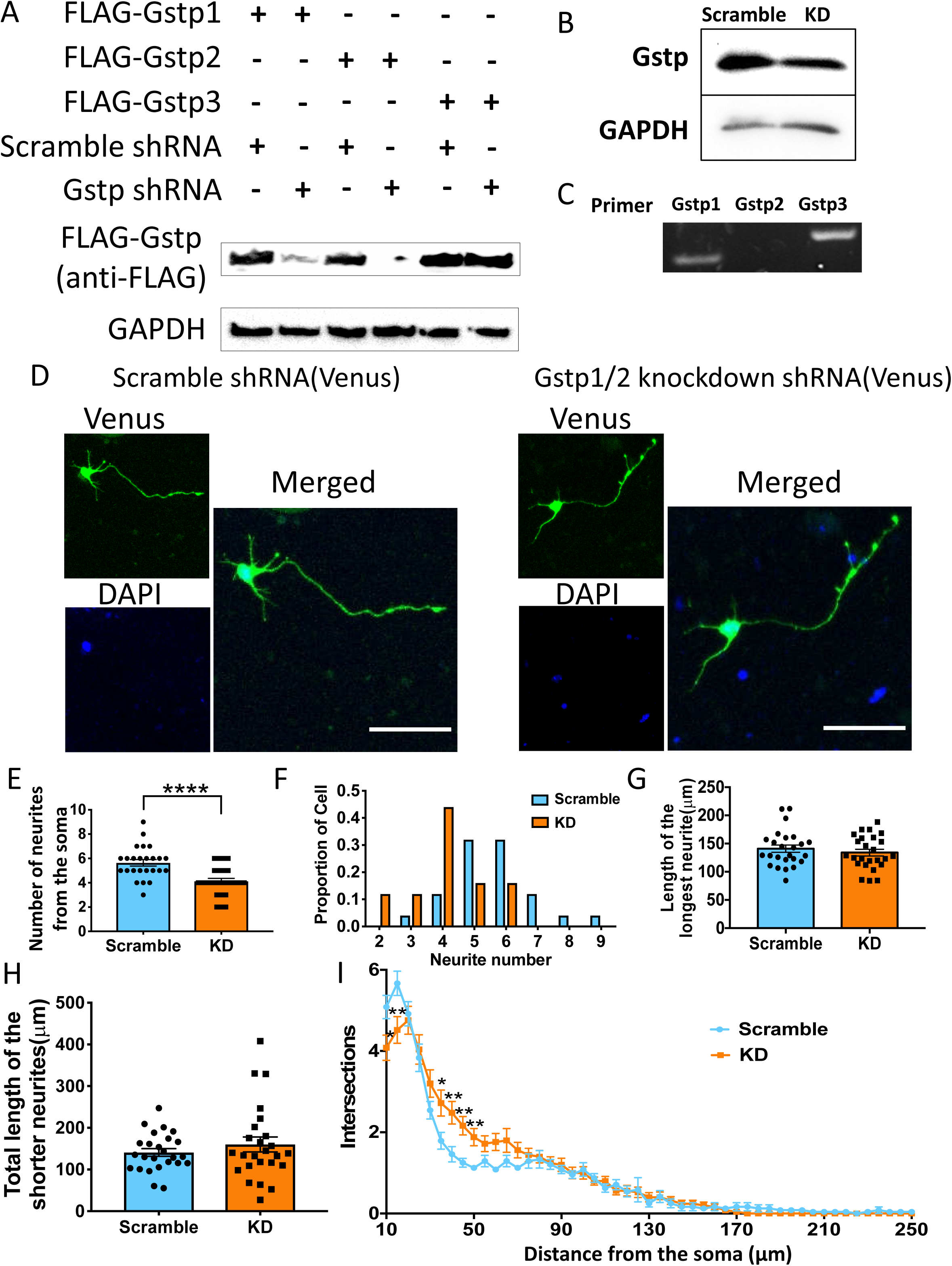
Knocking down of Gstp1 and 2 resulted in morphological defects in mouse primary cortical neurons. **A.** The plasmids encoding scramble or Gstp shRNA and FLAG-tagged Gstp1, 2 or 3 were co-transfected into HEK-293 cells. After 48 hours, knockdown efficiency of shRNA was evaluated by the WB using anti-FLAG antibody. shRNA can knock down both Gstp1 and 2 at the same time, but not Gstp3. **B.** Knockdown validation of Gstp shRNA in the endogenous Gstp expression in N-2a cells with anti-GSTP1 antibody. Note that the KD efficiency by Gstp shRNA was 41%. **C.** The analysis of the expression of Gstp1, 2 and 3 in N-2a cells by RT-PCR. N-2a cells express Gstp1 and 3, but not Gstp2. In N-2a cells Gstp1 an3 were expressed, but not gstp2. **D.** Representative photos of primary cortical neurons transfected with the plasmid coding scramble or Gstp shRNA, which also codes for Venus fluorescent protein. Scale bar, 50 μm. **E.** Quantification of the number of neurites from the soma in scramble and KD group. There is a significant reduction in the number of neurites in the KD group compared to the scramble group. N=25 per group, unpaired t-test, **** P<0.0001. **F.** Neurite count distribution of scramble and KD neurons. Most scramble neurons have 5-6 neurites, while KD neurons have 4. **G.** Quantification of the length of the longest neurite in scramble and KD neurons. There is no significant difference between them. **H.** Quantification of the length of shorter neurites in scramble and KD neurons. There is no significant difference between them. **I.** Sholl analysis showing the branching pattern of the scramble and KD neurons. There is a difference on the number of branches at 10 and 15 μm away from soma as well as 35 to 50 μm away from the soma. N=25 per group, unpaired t-test, * P<0.05, ** P<0.01.

To analyze the functions of Gstp1 and 2 in neuromorphogenesis, we transfected primary cortical neurons with the plasmid coding either the scramble shRNA or the Gstp shRNA and cultured for 48 hours as described in the methods (Figure 2D). We re-plated the neurons onto cover slips after 48 hours, which allowed time for the shRNA to be fully expressed before the re-plating occurred. Neurite length, neurite number, and branching pattern were analyzed at 48 hours after re-plating. We found the neurons transfected with KD plasmid (4.12 ± 0.2403) had less neurites at 48 hours after re-plating compared to the control neurons (5.64 ± 0.2638) transfected with the plasmid coding scramble shRNA (scramble, 5.64 ± 0.2638; KD, 4.12 ± 0.2403; unpaired t-test: t(48)=4.26, ****, p<0.0001) (Figures 2E). This suggests that Gstp proteins are essential for neurite initiation. Most of the cortical neurons have 5 to 6 neurites from the soma, while most of the KD neurons have 4 neurites from the soma (Figure 2F). No significant difference was observed in the length of the longest neurite, which will likely become the axon (scramble, 134 ± 5.94; KD 140.9 ± 6.424; unpaired t-test: t(4)=0.79, p=0.4312) (Figure 2G), as well as the length of the shorter neurites, which will likely become dendrites (scramble, 140.9 ± 9.114; KD, 160.1 ± 18.06; Unpaired t-test: t(4)=0.95, p=0.3485) (Figure 2H). Sholl analysis showed that the KD neurons had less neurite branching close to soma at 10 μm and 15 μm, but more branching from 35 μm to 50 μm (Figure 2I). These data suggest that Gstp1 and 2 are important for neurite initiation and neurite branching in a region-specific manner.

### Knockdown of Gstp1 and 2 in vivo showed abnormal dendrite morphology and defects in neurite initiation

To study the function of Gstp1 and 2 in neurite formation *in vivo*, we performed *in utero* electroporation (IUE) with the plasmids coding for scramble or Gstp1/2 KD shRNA. IUE was performed at E15.5 to mark pyramidal neurons in layers 2/3 (Brn2 positive) as previously described (Taniguchi et al., 2012). Gstp1/2 KD pyramidal neurons reached their final destination, layers 2/3 in the cortical plate, indicating that there were no defects in neurogenesis and neuronal migration (Figure 3A). Since *in vitro* knockdown experiments showed that Gstp1/2 KD neurons did not affect the length of the longest neurite, which likely becomes the axon, we focused the analysis on dendritic formation *in vivo*. At P3, we analyzed the morphology of neurons, which were visualized by Venus fluorescent proteins coded on the same plasmid coding for shRNA. We began our analysis at P3 so we could analyze the morphology of the apical dendrite, because basal dendrites have not emerged or are minimally emerged at this time point.

**Figure 3.**
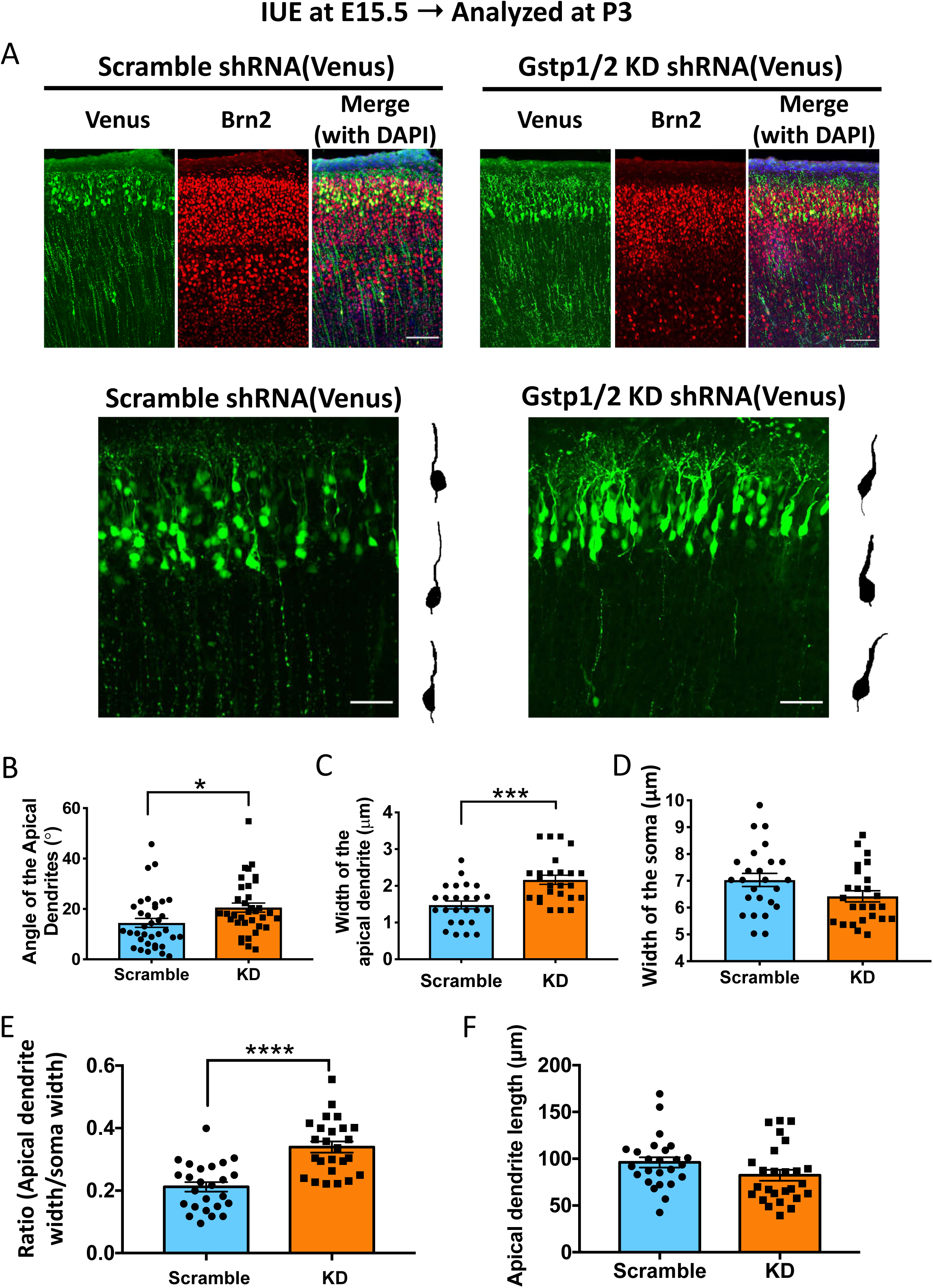
*In vivo* knockdown of Gstp1 and 2 using shRNA caused morphological defects in pyramidal neurons in layers 2/3 of the cerebral cortex at P3. **A.** Embryonic cortical neurons were electroporated with the plasmid coding for scramble or Gstp shRNA at E15.5 using *in utero* electroporation, and then the brain samples were analyzed at P3. Upper panel: *In utero* electroporation at E15.5 marked pyramidal neurons in layers 2/3 of the cortex. Brn2 is a marker for layers 2/3 and 5. Lower panels: Representative photos of Venus-positive pyramidal neurons electroporated with the plasmids coding for scramble shRNA (left panel) and Gstp shRNA (right panel). Scale bar 100 μm for the upper panels, scale bar 50 μm for the lower panels. **B.** Quantification of the angle of the apical dendrite in the scramble and KD neurons. Gstp KD caused significant increase of the angle of the apical dendrite. N=35 per group, unpaired t-test, * P<0.05. **C.** Quantification of the width of the apical dendrite. There is significant increase in the width of apical dendrite in KD neurons compared to scramble neurons. N=25 per group, unpaired t-test, *** P<0.001. **D.** Quantification of the width of the soma in control and KD neurons. There is no significant difference between them. **E.** The ratio of the width between the apical dendrite and the soma. There is significant increase in the ratio in KD neurons compared to scramble neurons. N=25 per group, unpaired t-test, **** P<0.0001. **F.** Quantification of the length of the apical dendrite in scramble and KD neurons. There is a tendency to the decrease of the length in KD neurons, but no significant difference in the length of apical dendrite in KD neurons compared to control ones. N=25 per group.

We measured the angle from the soma for the quantitative study of the apical dendrite orientation. For measuring the angle from the soma, we manually set up the reference line perpendicular to the pial surface of the brain. We found that KD neurons had increased angle of the apical dendrite compared to the scramble neurons (scramble, 14.48 ± 1.769 degrees; KD, 20.56 ± 1.822 degrees; unpaired t-test: t(68)=2.39, *, p=0.0194) (Figure 3B). We also observed a thicker apical dendrite in neurons deficient in Gstp1 and 2, especially in the proximal region of the apical dendrite (scramble, 1.480±0.1074 μm; KD, 2.166±0.125 μm; unpaired t-test: t(48)=4.166, ***, p<0.001) (Figures 3C). We measured the width of the soma both in the scramble and knockdown neurons, and there is no significant difference between the two groups (scramble, 7.032±0.2462 μm; KD, 6.423±0.2097 μm; unpaired t-test: t(48)=1.885, p=0.0655) (Figure 3D). Then, we calculated the ratio of the width of the apical dendrite against the width of the soma, and found that the width of the apical dendrite in KD neurons was significantly wider than in scramble neurons (scramble group, 0.2121±0.0151; knockdown group, 0.3394±0.01775; unpaired t-test: t(48)=5.464, ****, p<0.0001) (Figure 3E). Since there is no significant difference in the width of the soma between the control and KD neurons, the Gstp KD deficiency causes the swelling in the apical dendrite, not the entire cell. Taken together, these results suggest the importance of Gstp1/2 in proper morphogenesis of the apical dendrite. Meanwhile, there is no significant difference in the length of the apical dendrite between Gstp KD and scramble neurons, (scramble, 96.06±5.537 μm; KD, 82.21±6.192 μm; unpaired t-test: t(48)=1.67, p=0.1022) (Figure 3F). Thus, the Gstp KD caused the defects both in apical dendrite orientation and morphology, but not length.

### In vivo knockdown of Gstp1 and 2 using shRNA resulted in defects in neurite initiation of basal dendrites at P15

Next, we analyzed dendrite number, apical dendrite length, the width of the apical dendrite, and neurite branching at P15 (Figure 4). We confirmed that KD pyramidal neurons stayed in layers 2/3 (Brn2 positive) at P15 (Figures 4A). We found that knockdown of Gstp1 and 2 caused significant defect in neurite number from the soma compared to the scramble neurons (scramble, 4.92 ± 0.223; KD, 4.08 ± 0.199; unpaired t-test: t(48)=2.808, ** p=0.0072) (Figure 4B), reflecting the decrease of the number of basal dendrites, because we observed the correct formation of the apical dendrite in the KD neurons at P15. No defect was found in the length of the apical dendrite (scramble, 340.1 ± 26.68; KD, 292.5 ± 21.42; unpaired t-test: t(48)=1.393, p=0.1702) and the total length of the dendrites (scramble, 621.073 ± 56.448; KD, 491.443 ± 33.171; unpaired t-test: t(48)=1.98, p=0.0535). (Figure 4C and D). Although we found the abnormal width of the apical dendrite at P3, there is no difference in the width of apical dendrite, the soma, or the ratio between Gstp KD and control neurons at P15 (apical dendrite: scramble, 1.770 ± 0.111; KD, 1.646 ± 0.083; unpaired t-test: t(48)=0.89, p=0.377, soma: scramble, 11.66 ± 0.588; knockdown, 11.54 ± 0.29; unpaired t-test: t(48)=0.18, p=0.8555, Ratio: scramble, 0.1442 ± 0.0077; KD, 0.1561 ± 0.0087; unpaired t-test: t(48)=1.022, p=0.3117) (Figures 4E-G). Sholl analysis showed a decrease in intersections only close to soma when Gstp1 and 2 were knocked down, and there are no defects in branching pattern at distal regions (longer than 25 μm) from the soma (Figure 4H).

**Figure 4.**
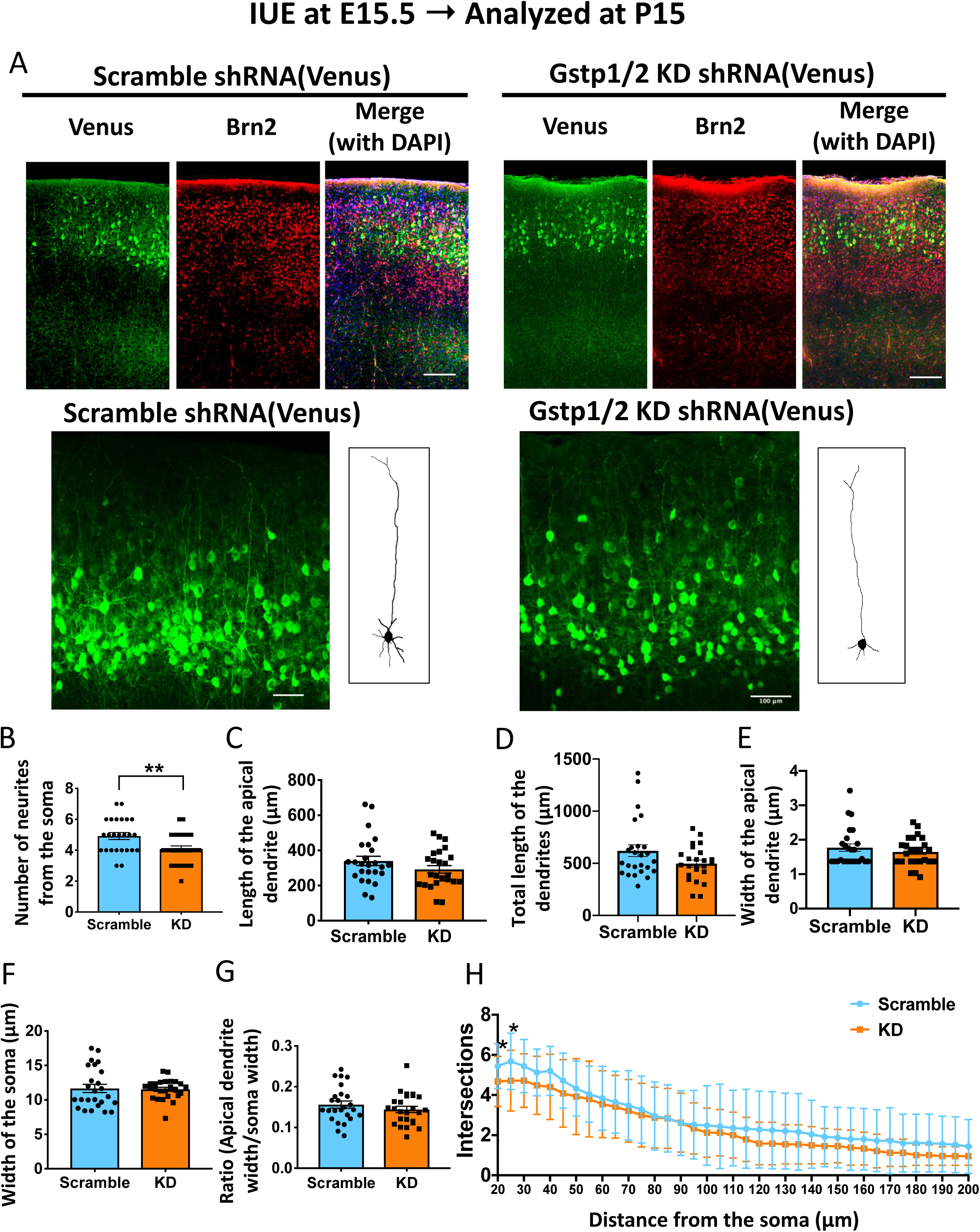
*In vivo* knockdown of Gstp1 and 2 using shRNA resulted in defects in neurite initiation at P15. **A.** Embryonic cortical neurons were electroporated with the plasmid encoding scramble or Gstp shRNA at E15.5 using *in utero* electroporation, and then brain samples were analyzed at P3. Upper panel: *In utero* electroporation at E15.5 marked pyramidal neurons in layers 2/3 of the cortex. Brn2 is a marker for layers 2/3 and 5. Lower panels: Representative photos of Venus-positive pyramidal neurons electroporated with the plasmids coding for scramble shRNA (left panel) and Gstp shRNA (right panel).Scale bar, 100 μm. **B.** Quantification of the number of neurites from the soma. The axon was excluded from the quantification. There is a significant reduction of the number of neurites in the KD neurons compared to the control neurons. N=25 per group, unpaired t-test. ** P<0.01. **C.** Quantification of the length of the apical dendrite in control and KD neurons. There is no difference in the length of the apical dendrite. **D.** Quantification of the total length of the dendrites. There is no difference in the total dendritic length. N=25 per group. **E.** Quantification of the width of the apical dendrite in the control and KD neurons. There is no difference in the dendritic width. N=25 per group. **F.** Quantification of the width of the soma in the control and KD neurons. There is no difference in the soma width. N=25 per group. **G.** Quantification of the ratio of the apical dendritic width vs. soma. There is no difference in the ratio between control and KD neurons. **H.** Sholl analysis showing the branching pattern of the control and KD neurons. There is a significant difference in branching at 20 μm and 30 μm away from the center of the soma. * P<0.05, N=25 per group.

Together, these results suggest that Gstp 1 and 2 are essential for proper morphogenesis of the apical dendrite at the early stage of neurite formation and the proper initial formation of basal dendrites at the later stage.

### Time-lapse live imaging revealed the importance of Gstp1 and 2 in the orientation of neurites in neurite initiation

We found that Gstp1 and 2 are essential for neurite initiation *in vitro* and *in vivo* (Figures 2 and 4). To study how Gstp KD affects the dynamics of neurite initiation, we utilized *ex vivo* time-lapse live imaging in combination with IUE (Figure 5A and B and Supplemental Videos 1 and 2). Mouse brain slices were prepared at P0 after IUE which was performed at E15.5, and then the process of neurite formation was recorded for 10 hours with 10 minute intervals.

**Figure 5.**
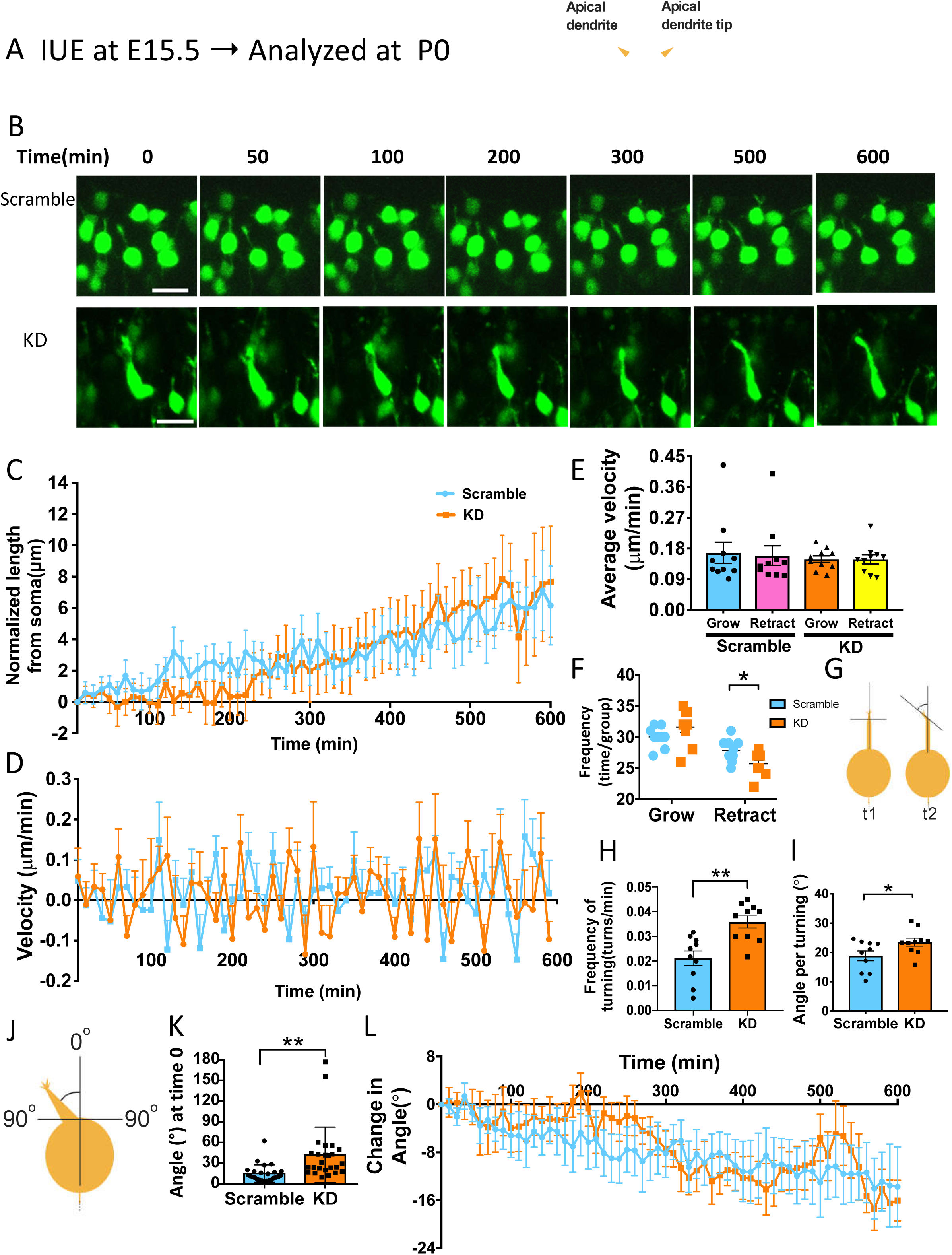
*Ex vivo* time-lapse live imaging at P0 showed defects in neurite orientation and dynamic movement of neurites in KD neurons. **A.** Cortical pyramidal neurons in the layers 2/3 were marked by electroporating with the plasmid coding scramble or KD shRNA at E15.5 using *in utero* electroporation, and then brain samples were analyzed at P0 by performing time-lapse live imaging on brain slices for a total period of 600 minutes with 10 minutes interval. **B.** Montage of the representative neurons electroporated with the plasmids coding for scramble or Gstp KD shRNA. **C** Normalized neurite length over time. The length of the neurite at each time point was measured for a time course of 600 minutes and normalized by subtracting the initial length of each neuron. N=10 per group. Scale bar, 20 μm. **D.** Measurement of growing or retracting velocity of neurite formation in scramble and KD neurons. **E.** Quantification of the average velocity of growth and retraction in the scramble and KD neurons. There is no significant difference in the neurite formation velocity in the scramble and KD group. **F.** Quantification of the frequency of growth and retraction in scramble and KD neurons. Note that there is no difference in the frequency of growth between scramble and KD group, but a significant difference in the frequency of retraction. N=10 per group, * P<0.05. **G.** Schematic of the measurement of tip turning. **H.** Quantification of the frequency of turning of the apical dendrite tip in the scramble and KD neurons. Apical dendritic tip of KD neurons turned more frequently than the scramble neurons. N=10 per group, ** P<0.01. **I.** Quantification of the angle per tip turning. The KD neurons made larger turning compared to the scramble neurons. N=10 per group, * P<0.05. **J.** Schematic of the measurement of the initial angle of the apical dendrite from the soma. **K.** Quantification of angle from the soma in the scramble and KD neurons at time point 0. N=20 per group, ** P<0.01. **L.** The change in angle of the apical dendrite over the time course of 600 minutes. N=10 per group.

At the beginning of the live imaging at P0, almost all pyramidal neurons arrived at the cortical plate and most of them started extending multiple neurites. In the wildtype condition, one of the neurites extends to the pial surface, and this likely becomes the apical dendrite. Another neurite often extends in the opposite direction toward the intermediate zone, and this likely becomes the axon. In addition to these two extensions, some neurons extend an additional couple of neurites from the lateral region of soma, but these neurites would disappear after repeating their extension and retraction several times, this happens at the early time points during the 10-hr live imaging from P0. In this study, all analyses were performed on the neurite likely to become the apical dendrite.

There was no significant difference in neurite length between the scramble and KD neurons at the beginning of the recording. To study the growth dynamics of the neurites in scramble and KD groups, the length at each time point was normalized by subtracting the length at time 0 from each (Figure 5C). The neurites on both scramble and KD neurons displayed constant growth and retraction throughout the time of recording, while the length of the apical dendrite in KD neurons was shorter at the beginning (0-250 minutes) (Figure 5C, orange line), it caught up to the same length as the control group after 250 minutes. During the neurite formation process, both scramble and KD neurons showed dynamic changes in the velocity of neurite formation(Figure 5D). There was no difference in the average velocity between the control and KD neurons (scramble grow, 0.673 ± 0.03116; scramble retract, 0.1589± 0.02867; KD grow, 0.1485 ± 0.009861; KD retract, 0.148 ± 0.01358) (Figure 5E). However, there is a significant difference in the frequency of retraction, where KD neurons had less retraction frequency during 10 hours than the scramble neurons, although there is no difference in growth frequency (scramble, 27.8 ± 0.5538; KD, 25.7 ± 0.5783; unpaired t-test: t(18)=2.623, *, p=0.0173) (Figure 5F). This could be the reason why the neurites on KD neurons catch up to the length of scramble neurites during live imaging, though KD neurites are shorter than the control at the early time points of the live imaging (Figure 5C).

During the recording, we noticed that neurite tips of KD neurons were more dynamically moving than the ones of the scramble neurons. Therefore, we analyzed the angle and frequency of the tip turning at each time point (Figure 5G). We found that the tip of the apical dendrite in the KD neurons turned more frequently than the control neurons during 10-hour recording (scramble, 12.7 ± 1.732; KD, 21.5 ± 1.47; unpaired t-test: t(18)=3.676, ** p=0.0011) (Figure 5H). We measured the angle every time the neurite changes direction (the average angle per turning of the apical dendrite tip) and found that KD neurons turned in larger angles than the scramble neurons (scramble, 18.83 ± 1.664 degrees; KD, 23.51 ± 1.352 degrees; unpaired t-test: t(18)=0.2096, * p=0.00424) (Figure 5I). Thus, the neurite tips in the KD neurons displayed more dynamic movement during the extension of the apical dendrite than the scramble neurons, indicating the role of Gstp1 and 2 in the outgrowth of the apical dendrite.

The angle of apical dendrite from soma was also analyzed (Figure 5J). The starting angles (time 0) were measured, and we found that the angle in the KD neurons was significantly larger than the scramble (scramble, 14.9 ± 2.529; KD, 42.16 ± 8.05; unpaired t-test: t(48)=3.23, ** p=0.0022) (Figure 5K). We also measured the angle every 10 minutes for 10 hours and calculated the changes in the angle (Figure 5L). A positive value represents an increase in the angle, in other words, the apical dendrite moves away from the perpendicular reference line and the pial surface. On the other hand, a negative value represents a decrease in the angle, indicating that the apical dendrite approaches the pial surface. We found that both the KD and scramble neurons decreased the angle over time, but the KD neurons more frequently changed the angle of the apical dendrite, indicating that KD neurons tried to rectify the angle to the level of the scramble group even though the apical dendrite in KD neurons emerged from the soma in the wrong direction.

Overall, these observations indicate that Gstp 1 and 2 are important for the formation of the apical dendrite orientation when the apical dendrite emerges from the soma, especially in the initial stage.

### Overexpression of Gstp1 and 2 in cortical neurons led to morphological defects

The knockdown experiments revealed the importance of Gstp1 and 2 in neuromorphogenesis *in vitro* and *in vivo* (Figures 2-5). Our shRNA knocked down both Gstp1 and 2 at the same time so that the function of each Gstp in neuromorphogenesis remains unrevealed. Although the overexpression is not a true functional analysis, it can still reveal important mechanistic information. Therefore, in order to study Gstp1 and 2 separately, we overexpressed Gstp1 or 2 in primary cortical neurons and co-transfected with plasmid encoding YFP to visualize the morphology (Figure 6A). Transfection was performed using primary cortical neurons prepared from E15.5 embryos, and then neurons were cultured for 48 hours followed by re-plating. We analyzed neurite number, neurite length, and branching pattern, by comparing with neurons transfected with YFP alone 48 hours following re-plating (control group). Overexpression of Gstp1 caused a significant decrease of total neurite length in comparison to YFP transfected neurons (control, 344.3±18.79 μm; Gstp1 overexpression, 244.9±11.72 μm; one-way ANOVA with Tukey’s multiple comparison, F_(2,72)_=1.989, * P<0.0001), as well as the length of shorter neurites (control, 186.7±14.89 μm; Gstp1 overexpression, 115.7±9.084 μm; one-way ANOVA with Tukey’s multiple comparison, F_(2,72)_=1.455, *** P=0.0006). There was no significant difference in the length of the longest neurite in Gstp1-overexpressing neurons (longest neurite length, 125.1±7.928 μm), compared to YFP transfected control neurons (longest neurite length, 152.1±9.625 μm; one-way ANOVA with Tukey’s multiple comparison, F_(2,72)_=1.097, ns, P=0.1390) (Figures 6B-D). The number of neurites from soma was also analyzed, but there was no significant difference between the control and Gstp1-overexpressing neurons (neurite number in the control neurons, 5.04±0.3978, neurite number in the KD neurons, 5.2±0.2449; one-way ANOVA with Tukey’s multiple comparison, F_(2,72)_=2.75, P>0.99) (Figure 6E). Sholl analysis showed that Gstp1 overexpression caused a significant decrease in neurite branching at proximal region from soma from 20 to 45μm (One-way ANOVA with Tukey’s multiple comparison, 20 μm, ** p=0.0132; 25 μm, *** P=0.0001; 30 μm, *** P=0.0006; 35 μm, *** P=0.0001; 40 μm, ** P=0.0093; 45 μm, * P=0.0308) (Figure 6F).

**Figure 6.**
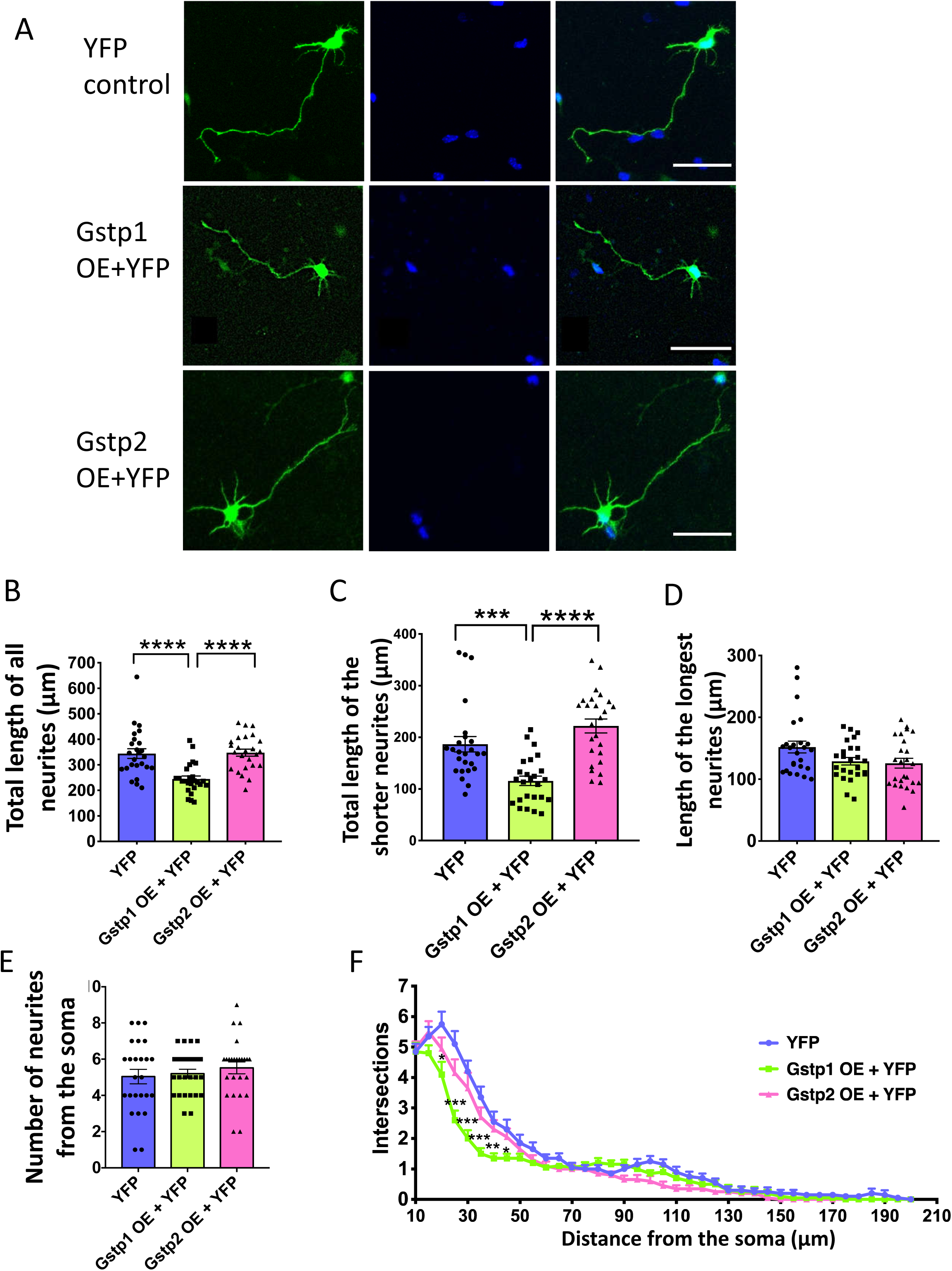
Overexpression of Gstp1 or 2 in primary cortical neurons suggests different functions in neurite formation. **A.** Representative photos of primary cortical neurons transfected with YFP alone (control), Gstp1 + YFP, or Gstp2 + YFP. Scale bar, 100 μm. **B.** Quantification of the total neurite length. One-way ANOVA suggests there was significant difference in the There is a significant reduction of total neurite length between the control and the Gstp1 OE groups. There is no significant difference between the control and Gstp2 OE groups. N=25 per group, one-way ANOVA for Tukey’s multiple comparison, F_(2,72)_=15.07, **** P<0.0001. **C.** Quantification of the sum of the shorter neurites’ length in control, Gstp1 OE, and Gstp2 OE neurons. There is a significant decrease between the control and the Gstp1 OE group. N=25 per group, one-way ANOVA for Tukey’s multiple comparison, F_(2,72)_=18.01, *** P<0.001. **D.** Quantification of the longest neurite’s length in control, Gstp1 OE, and Gstp2 OE neurons. No statistical significance was found. N=25 per group, one-way ANOVA for Tukey’s multiple comparison, F_(2,72)_=3.18. **E.** Quantification of the number of neurites from the soma. No significant difference was detected among the control, Gstp1 OE nor Gstp2 OE groups. N=25 per group, one-way ANOVA for Tukey’s multiple comparison, F_(2,72)_=0.6544, ns,p>0.05. **F.** Sholl analysis for quantifying branching pattern. Gstp1 OE neurons have fewer branches at 20 to 45 μm away from the soma. N=25 per group, one-way ANOVA for Tukey’s multiple comparison, * P<0.05, ** P<0.01, **** P<0.0001.

Overexpression of Gstp2 caused a slight but not significant increase in the length of shorter neurites (control, 186.7±18.42 μm; Gstp2 overexpression, 222 ± 13.25 μm; one-way ANOVA with Tukey’s multiple comparison, F_(2,72)_=1.455, P=0.1295) (Figure 6C, cobalt and pink bars). The length of the longest neurite length had a slight but not significant decrease (125.9±7.928 μm; one-way ANOVA with Tukey’s multiple comparison, F_(2,72)_=1.097, P=0.0708) (Figure 6D). No significant changes were observed in the length of the total neurites as well as neurite number from the soma (total neurite, 347.9 ± 13.81 μm, one-way ANOVA with Tukey’s multiple comparison, F_(2,72)_=1.989, p=0.9901; neurite number, 5.56 ± 0.3270, one-way ANOVA with Tukey’s multiple comparison, F_(2,72)_=2.75, P=0.5069) (Figures 6B and E). Sholl analysis showed a trend of increasing branching by Gstp2 overexpression, but it is not statistically significant (Figure 6F).

Overall, the overexpression experiments suggest that Gstp1 and 2 play important roles in neurite formation, and also that these two isoforms have different functions in neurite formation although they have high homology (Tables 1 and 2 and Figure 6).

### JNK specific inhibitor SP600125 rescued the decreased neurite number in Gstp1 and 2 KD primary cortical neurons

A previous study by Wang showed that GSTP1 directly interacts with JNK1 in the mouse embryonic fibroblast cell line, NIH3T3, and the enzymatic activity of JNK1 is inhibited by its interaction with GSTP1 (Wang et al., 2001). Also, Cdk5 activity is inhibited by Gstp (Sun et al., 2011). Previous studies have shown that JNK proteins and Cdk5 play important roles in neuritogenesis (Eom et al., 2005; Eminel et al., 2008). Therefore, we tested whether the inhibition of JNK activity can rescue the defects seen in Gstp KD primary cortical neurons. We applied SP600125, a specific JNK inhibitor which inhibits JNK1, 2, and 3 kinase activity with similar efficacy, in primary cortical neuron culture (Bennett et al., 2001) (Figure 7). SP600125 or DMSO (as a control) was added to the culture medium after 1 hour of re-plating before starting neurite initiation, which happens around 4 hours after re-plating (Dotti et al., 1988; Flynn, 2013). At 48 hours after re-plating, we fixed the cells and quantified the morphology by analyzing neurite number, because we have shown a fundamental function of Gstp1 and 2 is to regulate neurite initiation. Consistent with the Gstp KD neurons *in vitr*o and *in vivo* (Figures 2 and 4), the depletion of Gstp1 and 2 caused a significant reduction of neurite number with DMSO treatment (Figures 7A and B) (scramble and DMSO (brown bar), 5.36 ± 0.2880; KD and DMSO (orange bar), 4.28 ± 0.2347, two-way ANOVA with Tukey’s multiple comparison, F_(2,144)_=6.566, * P=0.0413). JNK inhibition with SP600125 rescued the defect in neurite number caused by Gstp knockdown as compared to control neurons (KD and DMSO (orange bar), 4.28 ± 0.2347; KD and SP600125 (green bar), 5.36 ±0.3208 two-way ANOVA with Tukey’s multiple comparison, F_(2,144)_=6.566, * P=0.0413) (Figure 7B). Although the importance of JNK in neurite formation is well known as described above, the treatment of neurons with 1µM SP600125 in scramble neurons did not show any defects in neurite number. However, we observed shorter neurite length in scramble neurons treated with 1µM SP600125, suggesting that 1µM is an effective concentration to inhibit JNK activity in neurite formation (data not shown). To ensure Gstp’s effects on neurite initiation are specifically through the JNK pathway and not through other related pathways known to affect neurite formation, we tested whether Cdk5 inhibition could also rescue these defects. Cdk5 activity is also negatively regulated by Gstp and is related to neurite formation (Nikolic et al., 1996; Paglini et al., 1998; Sun et al., 2011). The Cdk5 inhibitor Roscovitine was used to test whether it can also rescue the defects in neurite number in the KD neurons. By treating neurons transfected with the plasmid coding for scramble or Gstp shRNA with Roscovitine, we found that Cdk5 inhibition could not rescue the defects in neurite number in KD neurons (KD and DMSO (orange bar), 4.28 ± 0.2347: KD and Roscovitine (red bar), 436 ± 0.2227, two-way ANOVA with Tukey’s multiple comparison, ns, p>0.99; scramble and Roscovitine (blue bar), 5.48 ± 0.2318; KD and Roscovitine (red bar), 436 ± 0.2227, two-way ANOVA with Tukey’s multiple comparison, * P=0.0303; KD and DMSO (orange bar), 4.28 ± 0.2347: scramble and Roscovitine (blue bar), 5.48 ± 0.2318, two-way ANOVA with Tukey’s multiple comparison, * P=0.0158). We also tested a higher concentration (5 μM) of Roscovitine, but it still could not rescue the defects (data not shown).

**Figure 7.**
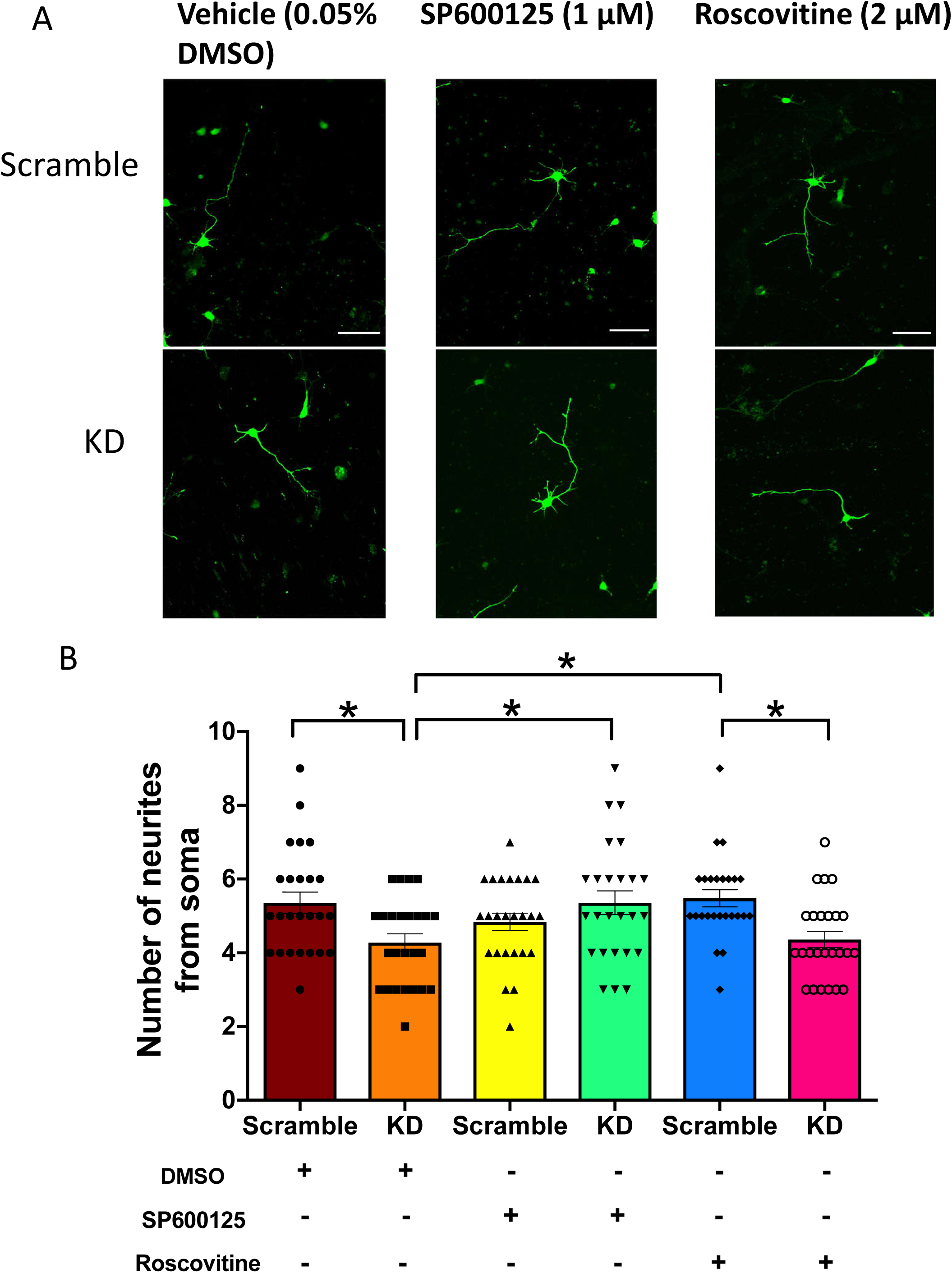
JNK inhibitor SP600125 rescued the defect in neurite initiation caused by Gstp1 and 2 knockdown. **A.** Representative photos of scramble or KD primary cortical neurons treated with vehicle,1 μM of SP600125, JNK inhibitor, or 2μM of Roscovitine, Cdk5 inhibitor. Scale bar, 20 μm. **B.** Quantification of the number of neurites from some in the scramble or KD neurons with or without the treatment of SP600125 or Roscovitine. The neurite number in KD neurons was rescued by the treatment of SP600125, but not Roscovitine. Two-way ANOVA determined that there was a statistical significance among the groups, N=25 per group, F(2,144)=6.566,* P<0.05.

Thus, the defects in neurite initiation caused by the knockdown of Gstp1 and 2 were rescued by the inhibition of JNK activity, but not Cdk5, indicating that Gstp proteins regulate neurite initiation specifically via the JNK signaling pathway.

## Discussion

In this study, we established that Gstp proteins are essential for neurite formation, particularly neurite initiation. As far as we know, this is the first study that elucidates the functions of Gstp proteins in neuritogenesis during cortical development. We showed that both overexpression and knockdown of the Gstp proteins resulted in defects in neuronal morphology during neurite formation. Knockdown of Gstp1 and 2 together using shRNA *in vitro* revealed a significant decrease in neurite number from the soma, indicating the importance of Gstp proteins in neurite initiation (Figure 2). Sholl analysis showed that neurite branching proximal to the soma was reduced in KD neurons, which resulted from defects in neurite initiation, whereas at 30 to 55 μm away from the soma, neurite branching was more frequent in KD neurons (Figure 2F). By *in vivo* KD of Gstp1 and 2, we found an abnormal swelling in the apical dendrite of KD pyramidal neurons in layer 2/3 at P3, while at P15, the swelling was not seen, indicating the defects were corrected as neurons matured (Figures 3E and 4G). Similarly, we observed a disrupted angle of the apical dendrite at P3, which was rectified and not seen at P15. At P15, we observed a reduced number of basal dendrites in KD pyramidal neurons, which is in chorus with our observations in primary cortical neurons *in vitro* (Figures 2E and 4B). This indicates that the regulation of neurite initiation is a primary function of Gstp1 and 2 in the developing cortex. Thus, Gstp1 and 2 are key players in neurite initiation and apical dendrite orientation at the early stage of neuritogenesis, but some of these deficiencies seem to be compensated for at later developmental stages. Live imaging at P0 showed that the length of extension of the apical dendrite in KD neurons was shorter than the control at the beginning of the live imaging, while the KD neurons increased the length of the apical dendrite after 250 minutes (Figure 5C). Although KD of Gstp1 and 2 did not affect the velocity of growth and retraction in the neurons (Figure 5E), by quantifying the frequency of growth and retraction of neurites, we found that the KD cells retracted neurites less frequently, indicating that this is the reason why the length of the apical dendrite caught up to the one of the scramble neurons after 250 minutes of the live imaging (Figure 5C and F). Also, we found the tip of the apical dendrite was more frequently and dramatically changing its direction in the KD neurons (Figures 5G-I). Thus, the live imaging revealed that the initial step of apical dendrite initiation was disrupted by Gstp 1/2 KD. Overexpression of Gstp 1 or 2 resulted in changes in neurite length, suggesting a role in neurite elongation (Figure 6). Finally, we found the defects in neurite initiation seen in KD neurons were rescued by the treatment of KD neurons with JNK inhibitor, SP600125 (Figure 7). This strongly indicates that the functions of Gstp proteins in neurite initiation are mediated by the JNK signaling pathway.

Humans have only one *GSTP, GSTP1*, whereas mice have three isoforms, *Gstp1, 2* and *3*. The human *GSTP1* gene has high homology to mouse *Gstp1* and *2* with about 83% identity, but less to mouse *Gstp3* with about 75% identity (Tables 1 and 2). Since both Gstp1 and 2 are highly homologous to each other and to human GSTP1, we analyzed the function of Gstp1 and 2 together. In this study, we used the shRNA by which Gstp1 and 2, but not Gstp3, were knocked down at the same time. The efficiency was approximately 85% for Gstp1, and 95% for Gstp2. Additionally, though there is a report indicating that Gstp is important for cell survival (Tew and Townsend, 2012), in our study the knockdown in primary neurons did not cause cell death within 5 days after shRNA transfection *in vitro*. Moreover, our *in vivo* experiments indicate that Gstp KD neurons were not eliminated *in vivo* for 19 days (P15) after IUE at E15.5, suggesting that the results obtained from Gstp1 and 2 knockdown here were not the secondary effects of an unhealthy condition.

Our analysis of the expression of each Gstp isoform during cortical development by using RT-PCR with the isoform-specific primers revealed that Gstp1, but not Gstp2 or 3, is the major Gstp protein expressed during embryonic cortical development, especially at the early stages (Figure 1E). To identify the specific roles of Gstp1 and 2 in neurite formation, we adopted the gain-of-function approach by overexpressing Gstp1 or 2 in primary cortical neurons (Figure 6). In our overexpression experiments, Gstp1 or Gstp2 overexpression caused different phenotypes in cortical neurons, suggesting the possibility that they have different roles in neuritogenesis despite their high similarity. Although humans have one GSTP, GSTP1, it is informative to clarify the function of Gstp1, 2, and 3 separately, and the information from these analyses will help us understand the mechanisms of neurite formation regulated by Gstp proteins in detail. To further analyze the functions of each Gstp protein, genomic modification by the CRISPR/Cas9 technique would be a useful tool. Also, the knockout mouse of each isoform would be useful to further test whether Gstp is important for neurobehavior. The Gstp1 and 2 double knockout mouse line has been created, and analysis of its phenotypes has been performed. An increase of skin tumorigenesis was observed (Henderson et al., 1998). Although the authors described that the double knockout mice appeared healthy with no defects in histopathology in major organs, it is unclear if the brain was analyzed. In addition, another *Gstp* knockout mouse line (*GstpΔ/Δ*) was created by deleting all three *Gstp* genes (Xiang et al., 2014). In the *GstpΔ/Δ* mice, no defects in the brain have been described, but it is not clear whether neurite formation was analyzed using these mice. Based on our observations and discussion here about the possibility that Gstp1 and 2 have different functions in neurite morphogenesis, the creation of the *Gstp1* and *Gstp2* single-gene knockout mice would be beneficial for understanding their functions in more detail.

*In vivo* knockdown of Gstp1 and 2 in layer 2/3 pyramidal neurons showed disrupted morphology at P0, P3, and P15 (Figures 3-5). At P0, when we were performing live imaging, we observed a disrupted angle in the apical dendrite of the KD neurons, while at P15, the angle became normal. Also, at P3, we saw an abnormal swelling at the stalk region of the apical dendrite of knockdown neurons, but this phenotype was not seen at P15. Thus, during cortical development, the swelling of the apical dendrite and its orientation defects were normalized. This could be a result of a compensatory effect by another Gstp, Gstp3. In RT-PCR, we saw that the expression of Gstp3 in the cortex starts later during cortical development and becomes higher at the neonatal stage (Figure 1E). Therefore, Gstp3 could compensate for Gstp1 and 2 depletion, resulting in alleviated phenotypes. In addition to the intrinsic compensation by Gstp3, exogenic compensation such as guidance cues and cell-cell interactions could help correct the defects, especially in orientation. This possibility is supported by the observation in our live imaging. In live imaging, the tip of the apical dendrite in KD neurons changed more frequently, like the tip was exploring the right direction.

As mentioned above, an interesting phenomenon found in our live imaging data is that the apical dendritic tips of the KD cells turned more than the control neurons, in terms of the frequency and the angle of turning (Figures 5G-I). It appeared as if the KD tips were exploring which path to go, while the normal tips were straight forward to one direction (Figures 3B and 5G-L). The growth of the neurite tip relies on the growth cone localized at the leading edge of the neurite. The receptors on the growth cone can receive guidance cues and transfer the signals inside the growth cone. In the movement of growth cones, F-actin and microtubules play essential roles, and the microtubules move into the filopodia along with the F-actin bundles. Then, more microtubules invade into the filopodia to extend. In contrast, filopodia that are not invaded by microtubules will be retracted (Lowery and Van Vactor, 2009). In our recording, the tip of KD neurons would pursue one direction at one-time point, then retract the filopodia, and extend other filopodia at another region of the growth cone, and this process would repeat again and again. Based on our observations, there could be some defects in the coupling of F-actin and microtubules occurring in the KD cells, where the stabilization of the filopodia is compromised. A previous study showed that this coupling process is largely dependent on microtubule plus-end complexes, F-actin retrograde flow, and the gradient of microtubule stabilizers and destabilizers (Rodriguez et al., 2003; Cammarata et al., 2016). It is consistent with our observations since the microtubule plus-end complexes (+TIPs), as well as the stabilizer and destabilizers, are heavily regulated by kinases, such as JNK. For example, the motor protein kinesin requires the binding of active JNK to regulate microtubule dynamics and +TIP CLIP-170 rescuing activity (Daire et al., 2009; Cammarata et al., 2016; Kawasaki et al., 2018). To further study the cellular mechanism, the markers of microtubule plus-end and F-actin can be used in live-imaging to visualize how these behave in the growth cone of the KD neurons.

Although the swollen apical dendrite recovered at P15, the increased width at the proximal region of apical dendrite was observed at P3 (Figures 3C-E). Swelling at the proximal region of the apical dendrite could be linked to the excess of the trafficking of intracellular organelles from soma. For example, the Golgi apparatus has been shown to accumulate at the base of the apical dendrite, and mitochondria are also trafficked towards the dendrites (Wu et al., 2015). It is possible that in the knockdown neurons, excessive organelles accumulate at the base of apical dendrites because effective outward organelle trafficking towards the distal area of the dendrite is prevented. These possibilities could be addressed by time-lapse live imaging using organelles’ markers.

In the present study, we also explored the potential involvement of JNK and Cdk5 in the Gstp signaling pathway in neuritogenesis by using JNK and Cdk5 inhibitors, because JNK and Cdk5 kinase activities are inhibited by GSTP1 (Figure 7) (Wang et al., 2001; Eminel et al., 2008; Sun et al., 2011). We conducted experiments with JNK inhibitor SP600125 and found that the treatment of Gstp KD neurons with SP600125 rescued the neurite initiation defects caused by Gstp1 and 2 knockdown. Applying Cdk5 inhibitor Roscovitine, however, did not rescue the decreased neurite number. Thus, the results suggest that the functions of Gstp in neurite formation are specifically mediated through the JNK pathway. Previous study showed that neurons with JNK2 and JNK3 knockout had defect in neurite initiation after 24 hours of culture, indicating that JNK2 and JNK3 are important for neurite initiation, but not JNK1(Barnat et al., 2010). Given that our shRNA knocks down Gstp1 and 2, Gstp1 and 2 would regulate neurite initiation through JNK2 and 3. To further study the molecular mechanism, a JNK specific knockdown approach could be used in combination with Gstp knockdown. Meanwhile, the further exploration of the downstream targets of JNK by comparing the phosphorylation status of the targets between control and KD neurons could help understand the Gstp/JNK signaling pathway in neuritogenesis in more detail.

## Acknowledgments

This work has been supported by a research grant from the NINDS (NS096098).

## References

Aaker JD, Elbaz B, Wu Y, Looney TJ, Zhang L, Lahn BT, Popko B (2016) Transcriptional Fingerprint of Hypomyelination in Zfp191null and Shiverer (Mbpshi) Mice. ASN neuro 8:1759091416670749.

Adler V, Yin Z, Fuchs SY, Benezra M, Rosario L, Tew KD, Pincus MR, Sardana M, Henderson CJ, Wolf CR, Davis RJ, Ronai Z (1999) Regulation of JNK signaling by GSTp. EMBO J 18:1321–1334.

Bakos J, Bacova Z, Grant SG, Castejon AM, Ostatnikova D (2015) Are Molecules Involved in Neuritogenesis and Axon Guidance Related to Autism Pathogenesis? NeuroMolecular Medicine 17:297–304.

Barnat M, Enslen H, Propst F, Davis RJ, Soares S, Nothias F (2010) Distinct roles of c-Jun N-terminal kinase isoforms in neurite initiation and elongation during axonal regeneration. J Neurosci 30:7804–7816.

Bennett BL, Sasaki DT, Murray BW, O’Leary EC, Sakata ST, Xu W, Leisten JC, Motiwala A, Pierce S, Satoh Y, Bhagwat SS, Manning AM, Anderson DW (2001) SP600125, an anthrapyrazolone inhibitor of Jun N-terminal kinase. Proc Natl Acad Sci U S A 98:13681–13686.

Bennison SA, Blazejewski SM, Smith TH, Toyo-Oka K (2020) Protein kinases: master regulators of neuritogenesis and therapeutic targets for axon regeneration. Cell Mol Life Sci 77:1511–1530.

Cammarata GM, Bearce EA, Lowery LA (2016) Cytoskeletal social networking in the growth cone: How +TIPs mediate microtubule-actin cross-linking to drive axon outgrowth and guidance. Cytoskeleton (Hoboken) 73:461–476.

Cornell B, Wachi T, Zhukarev V, Toyo-Oka K (2016) Regulation of neuronal morphogenesis by 14-3-3epsilon (Ywhae) via the microtubule binding protein, doublecortin. Hum Mol Genet 25:4405–4418.

Daire V, Giustiniani J, Leroy-Gori I, Quesnoit M, Drevensek S, Dimitrov A, Perez F, Pous C (2009) Kinesin-1 regulates microtubule dynamics via a c-Jun N-terminal kinase-dependent mechanism. J Biol Chem 284:31992–32001.

Darrow SM, Grados M, Sandor P, Hirschtritt ME, Illmann C, Osiecki L, Dion Y, King R, Pauls D, Budman CL, Cath DC, Greenberg E, Lyon GJ, McMahon WM, Lee PC, Delucchi KL, Scharf JM, Mathews CA (2017) Autism Spectrum Symptoms in a Tourette’s Disorder Sample. J Am Acad Child Adolesc Psychiatry 56:610-617.e611.

Diez-Roux G et al. (2011) A high-resolution anatomical atlas of the transcriptome in the mouse embryo. PLoS Biol 9:e1000582.

Dotti CG, Sullivan CA, Banker GA (1988) The establishment of polarity by hippocampal neurons in culture. J Neurosci 8:1454–1468.

Drubin DG, Feinstein SC, Shooter EM, Kirschner MW (1985) Nerve growth factor-induced neurite outgrowth in PC12 cells involves the coordinate induction of microtubule assembly and assembly-promoting factors. J Cell Biol 101:1799–1807.

Eminel S, Roemer L, Waetzig V, Herdegen T (2008) c-Jun N-terminal kinases trigger both degeneration and neurite outgrowth in primary hippocampal and cortical neurons. J Neurochem 104:957–969.

Eom D-S, Choi W-S, Ji S, Cho JW, Oh YJ (2005) Activation of c-Jun N-terminal kinase is required for neurite outgrowth of dopaminergic neuronal cells. NeuroReport 16:823–828.

Flynn KC (2013) The cytoskeleton and neurite initiation. Bioarchitecture 3:86–109.

Goto S, Kawakatsu M, Izumi S, Urata Y, Kageyama K, Ihara Y, Koji T, Kondo T (2009) Glutathione S-transferase pi localizes in mitochondria and protects against oxidative stress. Free Radic Biol Med 46:1392–1403.

Gupta S, Barrett T, Whitmarsh AJ, Cavanagh J, Sluss HK, Dérijard B, Davis RJ (1996) Selective interaction of JNK protein kinase isoforms with transcription factors. EMBO J 15:2760–2770.

Hand R, Polleux F (2011) Neurogenin2 regulates the initial axon guidance of cortical pyramidal neurons projecting medially to the corpus callosum. Neural Dev 6:30.

Harrill JA, Freudenrich TM, Machacek DW, Stice SL, Mundy WR (2010) Quantitative assessment of neurite outgrowth in human embryonic stem cell-derived hN2(tm) cells using automated high-content image analysis. NeuroToxicology 31:277–290.

Henderson CJ, McLaren AW, Moffat GJ, Bacon EJ, Wolf CR (1998) Pi-class glutathione S-transferase: regulation and function. Chem Biol Interact 111-112:69–82.

Kawasaki A, Okada M, Tamada A, Okuda S, Nozumi M, Ito Y, Kobayashi D, Yamasaki T, Yokoyama R, Shibata T, Nishina H, Yoshida Y, Fujii Y, Takeuchi K, Igarashi M (2018) Growth Cone Phosphoproteomics Reveals that GAP-43 Phosphorylated by JNK Is a Marker of Axon Growth and Regeneration. iScience 4:190–203.

Knight TR, Choudhuri S, Klaassen CD (2007) Constitutive mRNA Expression of Various Glutathione S-Transferase Isoforms in Different Tissues of Mice. Toxicol Sci 100:513–524.

Komulainen E, Zdrojewska J, Freemantle E, Mohammad H, Kulesskaya N, Deshpande P, Marchisella F, Mysore R, Hollos P, Michelsen KA, Magard M, Rauvala H, James P, Coffey ET (2014) JNK1 controls dendritic field size in L2/3 and L5 of the motor cortex, constrains soma size, and influences fine motor coordination. Front Cell Neurosci 8:272.

Lowery LA, Van Vactor D (2009) The trip of the tip: understanding the growth cone machinery. Nat Rev Mol Cell Biol 10:332–343.

Mannervik B, Alin P, Guthenberg C, Jensson H, Tahir MK, Warholm M, Jornvall H (1985) Identification of three classes of cytosolic glutathione transferase common to several mammalian species: correlation between structural data and enzymatic properties. Proc Natl Acad Sci U S A 82:7202–7206.

Monaco R, Friedman FK, Hyde MJ, Chen JM, Manolatus S, Adler V, Ronai Z, Koslosky W, Pincus MR (1999) Identification of a Glutathione-S-Transferase Effector Domain for Inhibition of jun Kinase, by Molecular Dynamics. J Protein Chem 18:859–866.

Nikolic M, Dudek H, Kwon YT, Ramos YF, Tsai LH (1996) The cdk5/p35 kinase is essential for neurite outgrowth during neuronal differentiation. Genes Dev 10:816–825.

Paglini G, Pigino G, Kunda P, Morfini G, Maccioni R, Quiroga S, Ferreira A, Caceres A (1998) Evidence for the participation of the neuron-specific CDK5 activator P35 during laminin-enhanced axonal growth. J Neurosci 18:9858–9869.

Perron JC, Bixby JL (1999) Distinct Neurite Outgrowth Signaling Pathways Converge on ERK Activation. Mol Cell Neurosci 13:362–378.

Pischedda F, Montani C, Obergasteiger J, Frapporti G, Corti C, Rosato Siri M, Volta M, Piccoli G (2018) Cryopreservation of Primary Mouse Neurons: The Benefit of Neurostore Cryoprotective Medium. Front Cell Neurosci 12:81.

Reese D, Drapeau P (1998) Neurite growth patterns leading to functional synapses in an identified embryonic neuron. J Neurosci 18:5652–5662.

Rodriguez OC, Schaefer AW, Mandato CA, Forscher P, Bement WM, Waterman-Storer CM (2003) Conserved microtubule-actin interactions in cell movement and morphogenesis. Nat Cell Biol 5:599–609.

Schaefer AW, Schoonderwoert VTG, Ji L, Mederios N, Danuser G, Forscher P (2008) Coordination of actin filament and microtubule dynamics during neurite outgrowth. Dev Cell 15:146–162.

Seow KH, Zhou L, Stephanopoulos G, Too H-P (2013) c-Jun N-terminal kinase in synergistic neurite outgrowth in PC12 cells mediated through P90RSK. BMC Neurosci 14:153.

Shen C-P, Chou IC, Liu H-P, Lee C-C, Tsai Y, Wu B-T, Hsu B-D, Lin W-Y, Tsai F-J (2014) Association of glutathione S-transferase P1 (GSTP1) polymorphism with Tourette syndrome in Taiwanese patients. Genet Test Mol Biomarkers 18:41–44.

Shen H-M, Liu Z-g (2006) JNK signaling pathway is a key modulator in cell death mediated by reactive oxygen and nitrogen species. Free Radic Biol Med 40:928–939.

Sun K-H, Chang K-H, Clawson S, Ghosh S, Mirzaei H, Regnier F, Shah K (2011) Glutathione-S-transferase P1 is a critical regulator of Cdk5 kinase activity. J Neurochem 118:902–914.

Taniguchi Y, Young-Pearse T, Sawa A, Kamiya A (2012) In utero electroporation as a tool for genetic manipulation in vivo to study psychiatric disorders: from genes to circuits and behaviors. Neuroscientist 18:169–179.

Tew KD, Townsend DM (2012) Glutathione-s-transferases as determinants of cell survival and death. Antioxid Redox Signal 17:1728–1737.

Thévenin AF, Zony CL, Bahnson BJ, Colman RF (2011) GST pi modulates JNK activity through a direct interaction with JNK substrate, ATF2. Protein Sci 20:834–848.

Toyo-oka K, Wachi T, Hunt RF, Baraban SC, Taya S, Ramshaw H, Kaibuchi K, Schwarz QP, Lopez AF, Wynshaw-Boris A (2014) 14-3-3ε and ζ regulate neurogenesis and differentiation of neuronal progenitor cells in the developing brain. J Neurosci 34:12168–12181.

Visel A, Thaller C, Eichele G (2004) GenePaint.org: an atlas of gene expression patterns in the mouse embryo. Nucleic Acids Res 32:D552–D556.

Wang T, Arifoglu P, Ronai Z, Tew KD (2001) Glutathione S-transferase P1-1 (GSTP1-1) inhibits c-Jun N-terminal kinase (JNK1) signaling through interaction with the C terminus. J Biol Chem 276:20999–21003.

Won H, Mah W, Kim E, Kim J-W, Hahm E-K, Kim M-H, Cho S, Kim J, Jang H, Cho S-C, Kim B-N, Shin M-S, Seo J, Jeong J, Choi S-Y, Kim D, Kang C, Kim E (2011) GIT1 is associated with ADHD in humans and ADHD-like behaviors in mice. Nat Med 17:566–572.

Wu YK, Fujishima K, Kengaku M (2015) Differentiation of Apical and Basal Dendrites in Pyramidal Cells and Granule Cells in Dissociated Hippocampal Cultures. PLOS ONE 10:e0118482.

Xiang Z, Snouwaert JN, Kovarova M, Nguyen M, Repenning PW, Latour AM, Cyphert JM, Koller BH (2014) Mice lacking three Loci encoding 14 glutathione transferase genes: a novel tool for assigning function to the GSTP, GSTM, and GSTT families. Drug Metab Dispos 42:1074–1083.

Yamauchi J, Miyamoto Y, Sanbe A, Tanoue A (2006) JNK phosphorylation of paxillin, acting through the Rac1 and Cdc42 signaling cascade, mediates neurite extension in N1E-115 cells. Exp Cell Res 312:2954–2961.

Zhang J, Grek C, Ye Z-W, Manevich Y, Tew KD, Townsend DM (2014) Pleiotropic functions of glutathione S-transferase P. Adv Cancer Res 122:143–175.

